# MtDNA heteroplasmy controls tumor immune reprogramming through mitochondrial translation

**DOI:** 10.64898/2026.01.31.693596

**Authors:** Ning Liu, Guoshi Chai, Qunyu Lv, Xiaoling Deng, Jiatong Chen, Qian Zhang, Leping Zhang, Xu Liu, Ruolan Tang, Zhe Xu, Shang Cai, Min Jiang

## Abstract

Mitochondrial DNA (mtDNA) mutations are among the most frequent genetic alterations in human cancers, yet their molecular mechanisms in shaping tumor progression and metastasis under physiological conditions remain poorly understood. Here, we show that mtDNA heteroplasmy acts as a tumor cell–intrinsic rheostat that orchestrates fate decisions between progression and suppression. Moderate heteroplasmy induces a ‘ribosomal storm’ of mitochondrial translation and ribosome biogenesis, driven by leucyl-tRNA synthetase 2 (LARS2). This response enhances metabolic fitness and licenses the secretion of S100A8/A9, enabling tumor cells to actively remodel the immune microenvironment through recruitment of myeloid-derived suppressor cells and exclusion of CD8⁺ T cells, thereby promoting immune evasion and metastatic progression. In contrast, high heteroplasmy impairs tumorigenesis due to oxidative phosphorylation collapse. Disruption of LARS2, mitochondrial translation, or S100A9 signalling abrogates tumor progression. Our findings establish mtDNA heteroplasmy as a key regulator of cancer evolution, linking mitochondrial genetic variation to immune modulation, and reveal the LARS2–mitochondrial translation–S100A8/A9 axis as a therapeutically targetable programme within the pro-tumorigenic window of mtDNA heteroplasmy.

## Introduction

Mitochondria are unique among mammalian cellular organelles in possessing their own genome ^1^—mitochondrial DNA (mtDNA)—which is highly prone to pathogenic mutations due to limited DNA repair capacity ^2^. Accumulating evidence indicates that mtDNA mutations are among the most prevalent genetic alterations in human cancers ^3–6^, arising from two distinct sources: somatic mutations acquired during an organism’s lifetime and inherited germline variants ^4^.

Although mtDNA mutations have been linked to diverse cancer phenotypes, their functional consequences remain context-dependent and often paradoxical. For example, specific complex I mutations drive the development of renal oncocytoma ^7^, whereas truncating mutations in *ND5*, another core subunit of complex I, are associated with improved therapeutic responses in melanoma ^8^ and better survival in colorectal cancer ^9^—highlighting the dual roles of mtDNA alterations in tumor biology.

A key contributor to this complexity is mtDNA heteroplasmy—the coexistence of mutant and wild-type mitochondrial genomes within a single cell ^10^. Unlike nuclear DNA mutations, which are typically binary (heterozygous or homozygous), mtDNA heteroplasmy exists on a continuous spectrum from 0% to 100%, introducing a non-Mendelian source of phenotypic variability that can influence tumor evolution ^11^. Moreover, heteroplasmy levels are dynamically regulated through genetic drift *in vivo*, adding an additional layer of complexity ^10^. However, most prior studies have relied on *in vitro* systems ^8,12,13^ or models with uncontrolled heteroplasmy ^14,15^, limiting mechanistic insights. As a result, the role of mtDNA heteroplasmy in tumor growth and metastatic progression remains poorly defined in physiologically relevant *in vivo* settings. Crucially, how varying heteroplasmy levels affect the tumor microenvironment—the primary target of modern cancer therapies—remains largely unknown, hindering the development of mitochondria-targeted therapeutic approaches. Here, we establish a genetically defined *in vivo* platform in which pathogenic mtDNA mutations in *ND5* (a core complex I subunit) and *TrnE* (encoding mitochondrial *tRNA^Glu^*) are maintained at precise heteroplasmy levels. This system enables the systematic dissection of mtDNA variant dosage effects on tumor evolution within an intact immune context. We demonstrate that moderate levels of mtDNA heteroplasmy in both mutant backgrounds create a pro-tumorigenic window, in which tumor growth and metastasis are promoted through a tumor cell-intrinsic mechanism involving LARS2-dependent activation of mitochondrial translation and subsequent secretion of S100A8/A9, collectively driving reprogramming of the tumor microenvironment toward an immunosuppressive state. In contrast, high heteroplasmy levels impair tumorigenesis due to the collapse of oxidative phosphorylation (OXPHOS).

Collectively, our findings establish mtDNA heteroplasmy as a critical regulator of cancer progression, acting at the interface between tumor cell metabolism and immune modulation. These results provide a conceptual framework for considering mitochondrial genetic variation as a non-nuclear determinant of tumor fate.

## Results

### MtDNA heteroplasmy exhibits a biphasic impact on tumor growth and metastasis

To characterize mtDNA mutations in breast cancer, we reanalyzed mtDNA mutation data from breast tumors in the Cancer Mitochondrial Atlas (TCMA) database ^3^. The majority of mtDNA variants were point mutations, including non-synonymous and silent mutations in protein-coding genes, tRNA/rRNA mutations, and D-loop mutation in non-coding regions (n=214 patients, Fig. 1a). In these breast cancer samples, C:G>T:A and T:A>C:G mutations were the most and second most common mutation types, respectively (Fig. 1a). Among the observed mtDNA mutations in breast cancer samples, variants in the D-loop region were the most prevalent. Of the 13 protein-coding genes, *COX1* and *ND5* were identified as the most frequently mutated genes in breast cancer (Fig. 1b). In addition to mRNA-coding mutations, we also detected a subset of transfer RNA (tRNA) mutations with elevated variant allele frequencies in cancer tissues, including mutations in *MT-TL1*, *MT-TM*, *MT-TH*, *MT-TQ*, *MT-TS1*, *MT-TN*, *MT-TS2*, *MT-TT*, *MT-TY*, and *MT-TE*.

**Fig. 1.**
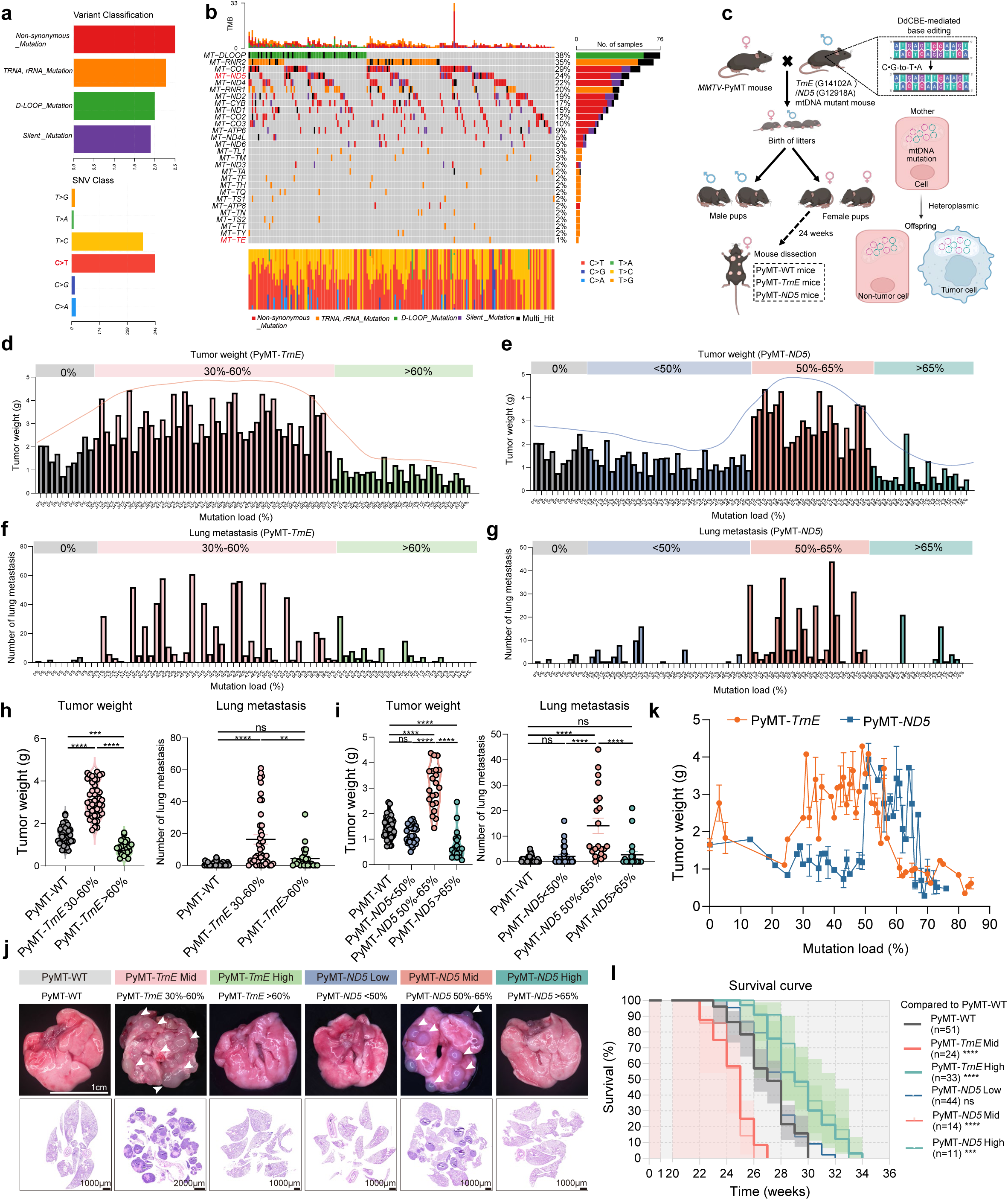
mtDNA heteroplasmy exhibits a biphasic effect on tumor progression in PyMT-*TrnE*/*ND5* breast cancer models. **a, b** Variant classification and SNV types of mtDNA mutations in breast cancer patients (n = 214) (**a**); Mutation oncoplot showing the proportion of mutations in different genes among coding and non-coding region (**b**). **c** Schematic showing the generation of an inherited mtDNA mutation model through mating of *MMTV*-PyMT mice with female *TrnE*/*ND5* mutant mice to produce PyMT-*TrnE*/*ND5* offspring. These mice develop spontaneous breast cancer carrying mtDNA mutations, enabling the study of mitochondrial genetic effects on tumorigenesis. **d, e** Tumor weight at different mutation load in PyMT-*TrnE* mice (**d**) and in PyMT-*ND5* mice (**e**). Trend lines were fitted using LOESS (Locally Estimated Scatterplot Smoothing) to illustrate the nonlinear relationship between mutation load and tumor weight. **f, g** Number of lung metastasis at different mutation load in PyMT-*TrnE* mice (**f**) and in PyMT-*ND5* mice (**g**). **h** Tumor weight (left) (PyMT-WT (n = 40), PyMT-*TrnE* 30%-60% (n = 43), PyMT-*TrnE* >60% (n = 25)) and number of lung metastasis (right) (PyMT-WT (n = 40), PyMT-*TrnE* 30%-60% (n = 41), PyMT-*TrnE* >60% (n = 23)). **i** Tumor weight (left) and number of lung metastasis (right) in PyMT-WT (n = 40), PyMT-*ND5*<50% (n = 29), PyMT-*ND5* 50%-65% (n = 21), PyMT-*ND5*>65% mice (n = 19). **j** Representative lung tissue images (top) (Scale bars, 1 cm) and H&E staining of lung sections (bottom) (Scale bars are shown in individual panels) showing the extent of metastasis in PyMT-WT, PyMT-*TrnE* 30%-60% (PyMT-*TrnE* Mid), PyMT-*TrnE* >60% (PyMT-*TrnE* High), PyMT-*ND5*<50% (PyMT-*ND5* Low), PyMT-*ND5* 50%-65% (PyMT-*ND5* Mid), PyMT-*ND5*>65% (PyMT-*ND5* High) mice. **k** Curves showing the relationship between tumor weight and mutation load in PyMT-*TrnE*/*ND5* mice. **l** Survival curves of PyMT-*TrnE*/*ND5* mice across Low, Mid, and High mtDNA mutation groups. n = 51, 24, 33, 44, 14 and 11 mice for PyMT-WT, PyMT-*TrnE* Mid/ High PyMT-*ND5* Low/Mid/ High group, respectively. Data are as mean ± s.e.m. One-way ANOVA with Tukey’s multiple comparison test (**h, i**), log-rank test (**l**). Schematic in **c** was created in BioRender. NS, not significant; \*\**P* < 0.01; \*\*\**P* < 0.001; \*\*\*\**P* < 0.0001.

To evaluate the potential impact of mtDNA mutations on breast tumor development, we bred the primary breast cancer mouse model (*MMTV*-PyMT) with female mutant *TrnE*/*ND5* mice. The mutant *ND5* and *TrnE* mice were generated using DddA-derived cytosine base editor (DdCBE) technology previously ^16–18^. The *ND5* mutation (m.12918G>A) mimics the human patient mutation at position 13513, while the *TrnE* mutation (m.14102G>A) represents a novel pathogenic mutation site corresponding to human position 14705. This breeding strategy enabled the generation of female offspring that develop breast cancer while carrying defined mtDNA mutations at variable heteroplasmy levels (referred to as PyMT-*TrnE* or PyMT-*ND5* models, respectively) (Fig. 1c). To verify that the tumors harbored only the intended single point mutations, mitochondrial whole-genome analysis of tumors confirmed high specificity for the induced mutations. Although low-frequency background mutations were detected, all were present at less than 5% frequency and were also observed in wild-type (WT) tumors (Extended Data Fig. 1a), thereby validating the fidelity of our models. We next assessed tumor size and lung metastasis in these mice at 24 weeks of age, when tumor development is typically apparent in PyMT models. We observed that mice with 30%-60% mtDNA mutation load in the PyMT-*TrnE* model and 50%-65% mutation load in the PyMT-*ND5* model exhibited significantly larger tumor sizes compared to PyMT-WT mice (Fig. 1d, e), along with increased numbers of lung metastatic nodules (Fig. 1f, g). These mice were classified as the moderate-mutant group. In contrast, the low-mutant group (PyMT-*ND5*<50%) showed tumor weights and lung metastasis counts comparable to those of PyMT-WT mice, while the high-mutant group (PyMT-*ND5*>65%, PyMT-*TrnE*>60%) exhibited smaller tumors and fewer lung metastases (Fig. 1h-j and Extended Data Fig. 1b). Histological analysis via H&E staining confirmed higher malignancy and increased metastatic foci in the moderate-mutant group compared to PyMT-WT, whereas reduced malignancy and fewer metastatic foci were observed in the high-mutant group (Fig. 1j and Extended Data Fig. 1b). The line graph further supported that moderate mutation levels correlate with larger tumor sizes (Fig. 1k). Consistent with enhanced proliferation, the moderate-mutant group showed stronger Ki67 staining, which was rare in the high-mutant group (Extended Data Fig. 1c). Survival analysis revealed shorter survival in the moderate-mutant group and prolonged survival in the high-mutant group relative to PyMT-WT (Fig. 1l). For simplicity, we refer to these groups as PyMT-WT, *TrnE*/*ND5*-Low, Mid, and High.

To further investigate how varying mtDNA mutation loads affect mitochondrial function in the tumors, we measured multiple mitochondrial protein levels in tumor tissues. Compared to controls, the levels of mitochondrial dynamics-related proteins (DRP1, MFN2) and GAPDH remained similar across all groups. However, mitochondrial DNA polymerase (POLG) was significantly upregulated in the PyMT-*TrnE*/*ND5* Mid group, and mitochondrial transcription factor A (TFAM) was significantly elevated in the PyMT-*ND5* Mid group (Extended Data Fig. 1d-f). Additionally, mitochondrial OXPHOS complex proteins (Complex I-V) and proteins involved in mitochondrial transport and protein folding (TOMM20, VDAC, HSP60) exhibited increased levels in the PyMT-*TrnE*/*ND5* Mid group, suggesting that moderate mtDNA mutations enhance mitochondrial biogenesis (Extended Data Fig. 1d-f). MtDNA copy number showed no significant differences among the groups (Extended Data Fig. 1g). Using transmission electron microscopy, we observed a significant increase in mitochondrial volume in tumor cells from the moderate-mutant groups (PyMT-*TrnE* Mid and PyMT-*ND5* Mid), although mitochondrial roundness remained unchanged (Extended Data Fig. 1h-j). Collectively, these findings demonstrate that moderate mtDNA mutations promote tumor growth and metastasis, accompanied by enhanced mitochondrial biogenesis.

### MtDNA mutation in tumor cells determines tumor growth and metastasis

To investigate the promotion of tumor growth associated with moderate mtDNA mutations, we analyzed the mtDNA mutation loads in tumors from established mouse models. We measured mutation loads in primary tumor tissues, metastatic tumor tissues, surrounding tissues (lung), and blood from PyMT-*TrnE*/*ND5* mouse models, normalizing against the mutation load in the tail. Our analysis revealed distinct mutation load patterns between the PyMT-*TrnE* and PyMT-*ND5* models. In the PyMT-*TrnE* Mid model, the mutation loads in randomly selected primary and lung metastatic tumor tissues exhibited variable patterns compared to the tail. However, in the PyMT-*TrnE* High group, mutation loads closely resembled those of the tail (Fig. 2a). In contrast, in the PyMT-*ND5* model, the mutation loads in tumor and lung metastatic tumor tissues from the Low, Mid, and High groups were generally lower than those in the tail (Fig. 2b). Across all groups, mutation loads in blood and lung tissues were comparable to those in the tail, indicating that tumor tissues display higher mutation heterogeneity relative to non-tumor tissues. To assess whether mutation load correlated with tumor size, we measured mutation loads in multiple tumors from the same mouse and observed a negative correlation between mutation load and tumor size (Fig. 2c). Notably, even tumors from the same mouse exhibited significant differences in mutation load (Fig. 2d). This variation was further evident when a single tumor was divided into three sections, each displaying distinct mutation loads (Fig. 2e). Larger tumors generally exhibited lower average mtDNA mutation loads.

**Fig. 2.**
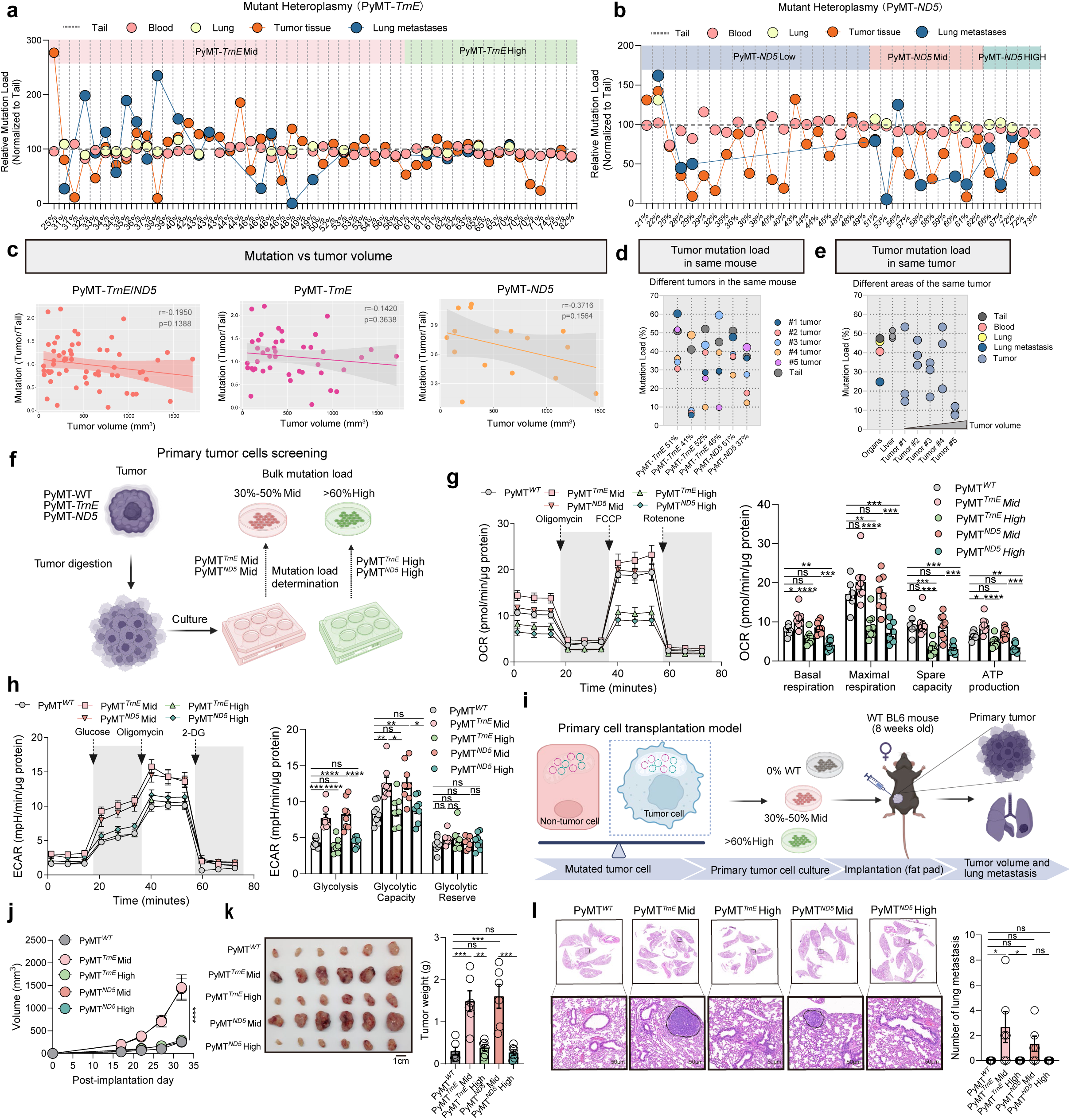
Moderate mtDNA mutation in primary tumor cells drives enhanced tumor growth and metastasis through cell-intrinsic metabolic reprogramming. **a, b** mtDNA mutation load in tail, blood, lung, primary tumor, and metastatic lung lesions from PyMT-*TrnE* (**a**) and PyMT-*ND5* (**b**) mice with varying mutation levels. **c** Linear regression analysis showing the correlation between tumor volume and mtDNA mutation load in PyMT-*TrnE* and PyMT-*ND5* mice. **d** mtDNA mutation load analysis of tumor tissues from different mammary glands within the same mouse. **e** mtDNA mutation load analysis of triplicate sections from the same primary tumor, along with matched blood, lung, metastatic lung tumor, and liver tissues (as control). **f** The strategy of the primary tumor cell isolation based on mutation load from PyMT-WT and PyMT-*TrnE*/*ND5* mouse models. Tumor cells with 30-50% mtDNA mutation level were classified as the moderate-mutant group, and those with >60% mutation level were assigned to the high-mutant group. **g** Oxygen consumption rate (OCR) (left) and quantification (right) of basal respiration, maximal respiration, spare capacity, ATP production in PyMT primary tumor cells with different mtDNA mutation loads. n = 6-8 replicates per group. **h** Extracellular acidification rate (ECAR) (left) and quantification (right) of glycolysis, glycolytic capacity, glycolytic reserve in PyMT primary tumor cells with different mtDNA mutation loads. n = 6-8 replicates per group. **i** Schematic of the orthotopic mammary engraftment strategy for primary tumor cells derived from PyMT*^WT^*, and PyMT*^TrnE^* Mid/High and PyMT*^ND5^* Mid/High groups. **j, k** Growth curves **(j)** and tumor weights **(k)** after dissection of primary tumor cells derived from PyMT*^WT^*, and PyMT*^TrnE^* Mid/High and PyMT*^ND5^* Mid/High groups (n = 6 mice per group). Scale bars, 1 cm. **l** Representative images of lung metastases (left) and quantification of metastatic nodules (right) following orthotopic implantation of primary tumor cells from PyMT*^WT^*, and PyMT*^TrnE^* Mid/High and PyMT*^ND5^* Mid/High groups (n = 6 mice per group). Scale bars, 50 μm. Data are as mean ± s.e.m. two-way ANOVA (**j**) and One-way ANOVA with Tukey’s multiple comparison test (**g, h, k** (tumor weight)**, l**). Schematic in **f** and **i** were created in BioRender. NS, not significant; \**P* < 0.05; \*\**P* < 0.01; \*\*\**P* < 0.001; \*\*\*\**P* < 0.0001.

We next employed mtscATAC-seq ^19,20^ to examine mutation loads at the single-cell level within tumor tissues. Our analysis identified eight distinct subpopulations (Extended Data Fig. 2a-c), with tumor cells being the most abundant (Extended Data Fig. 2d). Single-cell mutation load analysis revealed substantial heterogeneity across all cell types, with mutation loads ranging from 0% to 100% (Extended Data Fig. 2e). While the mean mutation load in non-tumor cells from all groups was close to that of their tails, tumor cells exhibited variable mutation loads (Extended Data Fig. 2e), underscoring the intrinsic complexity of mtDNA heteroplasmy in tumor cells. These findings indicate that mtDNA heteroplasmy levels differ markedly among tumor and non-tumor cells at the single-cell level, with tumor cells exhibiting greater heterogeneity than other cell types.

Furthermore, GO pathway analysis of significantly enriched pathways derived from open chromatin regions in tumor cells—based on mtscATAC-seq data comparing the Mid and High mutation groups—revealed marked upregulation of pathways associated with cell proliferation and migration (Extended Data Fig. 2f). Notably, genes linked to “regulation of myeloid cell differentiation” and “positive regulation of myeloid leukocyte differentiation” were also enriched in tumor chromatin accessibility profiles, suggesting that tumor cells with moderate mtDNA mutations may actively shape the immune microenvironment by potentiating myeloid cell differentiation (Extended Data Fig. 2f).

To determine whether mutant tumor cells are the primary drivers of enhanced tumor growth, we isolated primary tumor cells from PyMT-WT, *TrnE/ND5*-Low, Mid, and High groups to compare their proliferative capacities (Fig. 2f). We used control primary tumor cells (PyMT*^WT^*) from the PyMT-WT group, moderate-mutant primary tumor cells (PyMT*^TrnE/ND5^*Mid) with mutation loads of 30%-50% from the PyMT-*TrnE*/*ND5* Mid group and high-mutant primary tumor cells (PyMT*^TrnE/ND5^*High) with mutation loads exceeding 60% from the PyMT-*TrnE*/*ND5* High group for subsequent experiments (Fig. 2f). To assess mitochondrial and glycolytic activity, we measured oxygen consumption rate (OCR) and extracellular acidification rate (ECAR). Moderate-mutant tumor cells exhibited a modest increase in OCR but a significant elevation in ECAR compared to high-mutant cells, indicating a shift toward enhanced glycolytic metabolism. In contrast, high-mutant cells displayed significantly reduced OCR, reflecting impaired mitochondrial respiration, while ECAR levels remained largely unchanged (Fig. 2g, h). To evaluate the *in vivo* growth and metastatic potential of these cells, we inoculated primary tumor cells from different groups into WT recipient mice (Fig. 2i). Tumors derived from transplanted moderate-mutant primary tumor cells exhibited significantly enhanced proliferation and metastasis, while tumors derived from transplanted high-mutant primary tumor cells showed growth rates similar to control cells (Fig. 2j-l). Thus, we conclude that moderate-mutant tumor cells are the primary drivers of enhanced tumor growth.

To test whether the pro-tumorigenic and pro-metastatic effects associated with moderate mtDNA mutation loads are generalizable across different tumor types, we utilized the highly metastatic 4T1 murine breast cancer cell line. We performed direct mtDNA base editing *in vitro* to generate isogenic cell lines with defined mutation loads, followed by orthotopic transplantation into WT mice—thereby eliminating confounding effects of a mutant tumor microenvironment. Based on mutation load, we categorized these cell lines as 4T1-*TrnE*/*ND5* Low (0%-10%), Mid (40%-60%), and High (>80%) (Fig. 3a). OXPHOS protein expression was significantly upregulated in the moderate-mutant group, particularly for complex II, compared to the low-mutant group (Fig. 3b). Upon transplantation into WT recipient mice (Fig. 3c), moderate-mutant 4T1 cells formed larger primary tumors and exhibited a significant increase in lung metastasis compared to the low-mutant group, whereas high-mutant cells did not display this phenotype (Fig. 3d-h). Notably, tumors derived from moderate-mutant 4T1 cells displayed markedly expanded necrotic regions (Fig. 3i-k), further highlighting the aggressive *in vivo* behavior associated with moderate mtDNA mutation loads.

**Fig. 3.**
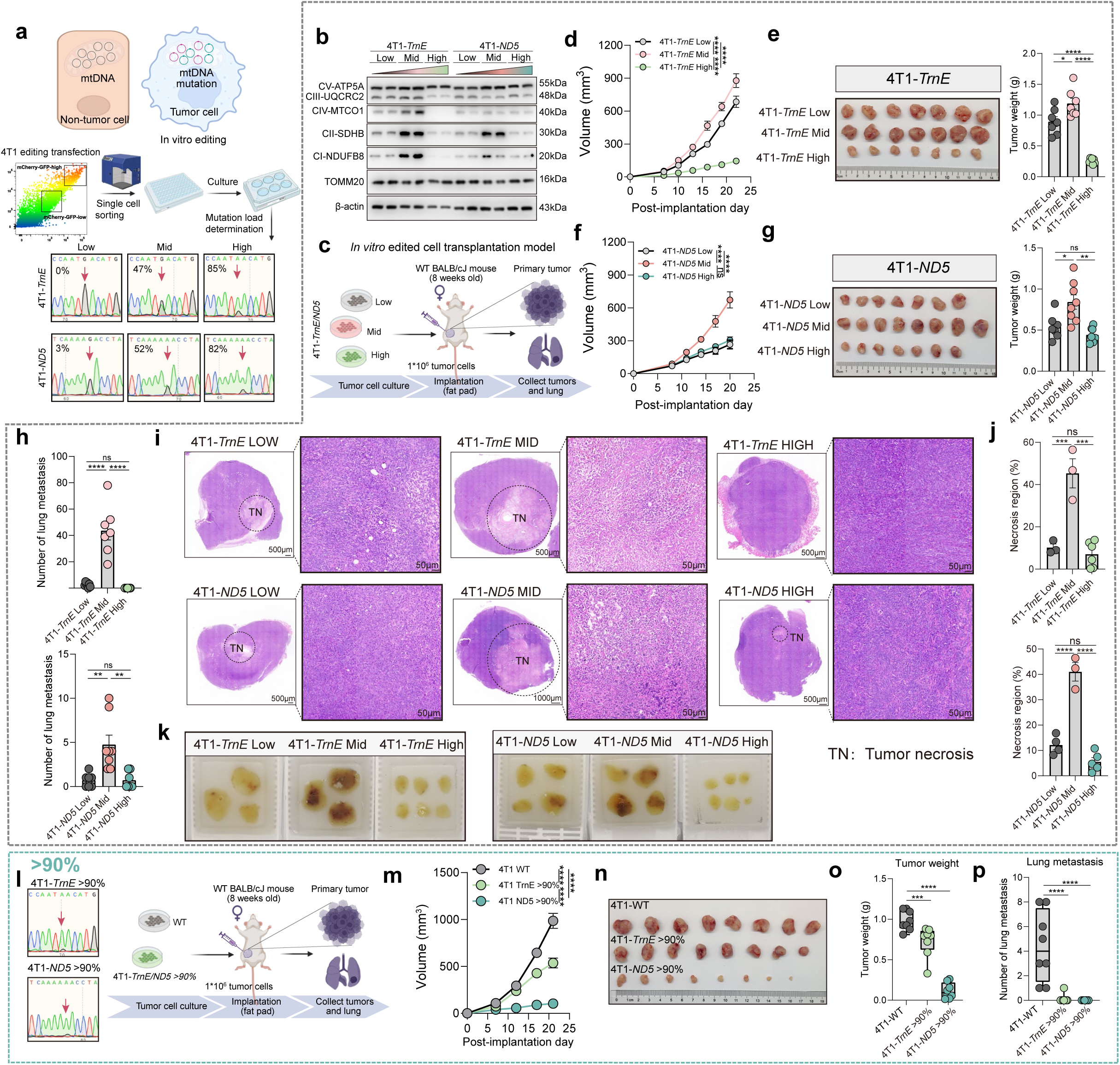
MtDNA mutation load modulates tumor growth and metastasis in the 4T1 transplantation model. **a** Schematic of the mtDNA editing strategy in 4T1 cells. 4T1 cells were transfected with DdCBE plasmids to introduce mutations at the m.14102 and m.12918 sites. Following single-cell cloning, clones were expanded and subjected to mutation load analysis. Based on the heteroplasmy levels, clones were classified into 4T1-*TrnE*/*ND5* Low, Mid, and High groups for subsequent functional studies. **b** Immunoblots showing expression levels of mitochondrial Complex I-V and TOMM20 in 4T1-*TrnE*/*ND5* Low, Mid, and High cells. n = 2 independent clones per group. **c** Schematic of the tumor cell engraftment strategy, showing the injection of 4T1-*TrnE*/*ND5* Low, Mid, and High cells into wild-type (WT) BALB/cJ mice for *in vivo* tumor growth and metastasis studies. **d, e** Growth curves of 4T1-*TrnE* Low, Mid, and High tumors after orthotopic implantation **(d)**. Representative tumor images (left) and quantification of tumor weights (right) are shown post-dissection **(e)** (n = 7 mice per group). **f, g** Growth curves of 4T1-*ND5* Low, Mid, and High tumors after orthotopic implantation **(f)**. Representative tumor images (left) and quantification of tumor weights (right) are shown post-dissection **(g)** (n = 7-8 mice per group). **h** Quantification of lung metastasis in 4T1-*TrnE/ ND5* Low, Mid, and High groups (n = 7 mice per group). **i-k** Representative H&E images of tumors from 4T1-*TrnE*/*ND5* Low, Mid, and High groups **(i)** (Scale bars are shown in individual panels) along with quantification of necrotic area **(j)** (n = 3-6 tumors per group). Photographs showing the gross morphology of tumors after dehydration and embedding **(k)**. **l** Schematic of the tumor cell engraftment strategy, showing the injection of 4T1-WT and 4T1-*TrnE*/*ND5* >90% cells into wild-type (WT) BALB/cJ mice. **m-o** Growth curves **(m)** of 4T1-WT and 4T1-*TrnE*/*ND5* >90% tumors after orthotopic implantation. Representative tumor images **(n)** and quantification of tumor weights **(o)** are shown post-dissection (n = 8 mice per group). **p** Quantification of lung metastases following orthotopic implantation of 4T1-WT and 4T1-*TrnE*/*ND5* >90% tumor cells. Metastatic nodules were counted post-dissection (n = 8 mice per group). Data are as mean ± s.e.m. two-way ANOVA (**d, f, m**) and One-way ANOVA with Tukey’s multiple comparison test (**e, g, h, j, o, p**). Schematic in **a**, **c** and **l** were created in BioRender. NS, not significant; \**P* < 0.05; \*\**P* < 0.01; \*\*\**P* < 0.001; \*\*\*\**P* < 0.0001.

Given that moderate mtDNA mutation loads accelerate tumor growth, the phenotypic impact of high mutation burdens may be masked in standard observation windows. To further evaluate the impact of high-level mtDNA mutations on tumorigenesis, we analyzed a subset of edited tumor cells with >90% mutation load (Fig. 3l). These 4T1 cells with >90% mutation load (*TrnE* or *ND5*) showed markedly reduced tumor growth (Fig. 3m-o) and fewer metastatic foci (Fig. 3p) upon orthotopic transplantation compared to WT controls. This suppression of tumor progression at very high mutation loads contrasts with the aggressive phenotype observed at moderate mutant level, supporting a biphasic, threshold-dependent role of mtDNA mutation burden in shaping tumor fate.

To determine whether mtDNA mutations in host-derived cells — particularly immune cells—contribute to tumor progression (Extended Data Fig. 3a), we quantified mtDNA mutation loads in sorted tumor-infiltrating immune populations. Mutation levels in neutrophils, macrophages, T cells, B cells, and NK cells were slightly reduced (except for B cells in PyMT-*ND5* tumors), suggesting purifying selection (Extended Data Fig. 3b). To assess whether *TrnE/ND5* mutant host environments influence tumorigenesis, we transplanted PyMT-WT tumor cells into *TrnE/ND5* mutant mice with varying mutation loads. Neither moderate-mutant nor high-mutant *TrnE/ND5* recipient mice showed significant differences in tumor growth compared to WT controls (Extended Data Fig. 3c-e). Further immunophenotyping of blood and tumor tissues revealed no significant differences in the proportions of macrophages, MDSCs, or T cells among recipient groups (Extended Data Fig. 3f, g).

Collectively, these findings demonstrate that moderate levels of mtDNA mutations in tumor cells—not in the surrounding tumor microenvironment—enhance tumor growth and metastatic potential, whereas high-mutant cells exhibit reduced tumor growth due to impaired mitochondrial function.

### Ribosomal storm-driven activation of mitochondrial translation promotes the growth of moderate-mutant tumor cells

To uncover intrinsic mechanisms underlying the fitness advantage of moderate-mutant tumor cells, we performed proteomic profiling of spontaneous tumors from PyMT-WT/*TrnE*/*ND5* mice. Proteomic PCA analysis showed a mild distinction between PyMT-*TrnE*/*ND5* Mid and High tumors (Fig. 4a). In PyMT-*TrnE*/*ND5* Mid tumors, approximately 30% of the detected mitochondrial proteins (MitoCarta3.0) exhibited significant changes, compared to only about 10% in PyMT-*TrnE*/*ND5* High tumors. Moreover, over 20% of mitochondrial proteins were significantly upregulated in the moderate-mutant group compared to the PyMT-WT group (Fig. 4b). Both mtDNA- and nuclear-encoded OXPHOS subunits displayed an increasing trend across all groups, with the most pronounced elevation in PyMT-*TrnE* Mid tumors (Fig. 4c). Tumors from moderate-mutant groups showed significant upregulation of pathways associated with mitochondrial translation (e.g., Mrpl, Mrps), mitochondrial gene expression, and OXPHOS (e.g., Cox5a, Coq9, Atp5pf, Shmt2, Tafazzin, Ndufb2, Ndufab1, Uqcrh) (Fig. 4d). In contrast, gene set enrichment analysis revealed significant downregulation of pathways involved in antigen processing and presentation (Fig. 4e).

**Fig. 4.**
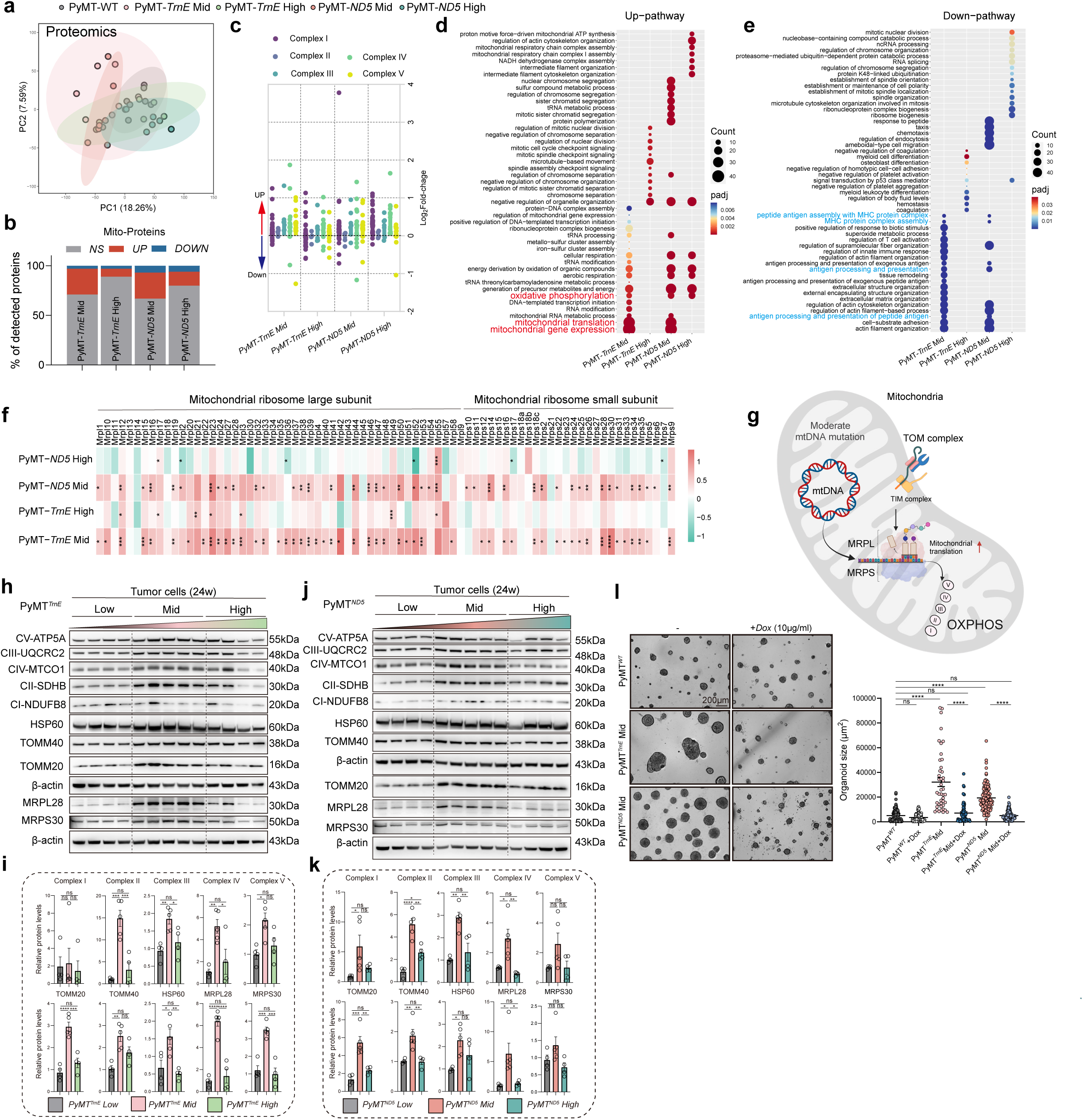
Moderate mtDNA heteroplasmy enhances mitochondrial translation and ribosomal biogenesis in a mutation type–independent manner. **a** Proteomics analysis of PyMT-WT, PyMT-*TrnE* Mid/High PyMT-*ND5* Mid/High tumors visualized by PCA. Each circle represents one mouse. n = 7 mice per group. **b** MitoCarta3.0 classification of detected proteins based on mitochondrial localization. **c** Respiratory chain subunits (Complex I-V) change from proteomics. **d, e** GO pathway enrichment analysis based on proteomics data from PyMT-*TrnE* Mid/High and PyMT-*ND5* Mid/High tumors compared to PyMT-WT. Upregulated **(d)** and downregulated pathways **(e)** are shown as a bubble plot, where bubble size indicates the number of enriched proteins (count), and color intensity reflects the adjusted p-value (padj). **f** Heatmap of log_2_ fold-change of MRPL and MRPS mitochondrial ribosomal proteins in bulk tumor proteomes from PyMT-WT, PyMT-*TrnE*, and PyMT-*ND5* groups (n = 7 mice per group). *P* values from t-test vs PyMT-WT. **g** Schematic showing mtDNA-mediated translation via MRPL and MRPS ribosomal subunits. The TOM complex facilitates the import and outer membrane localization of mitochondrial proteins, while inner membrane-localized respiratory complex subunits are influenced by mitochondrial translation efficiency. **h, i** Immunoblots **(h)** showing expression levels of mitochondrial OXPHOS proteins, mitochondrial transport and folding proteins, and mitochondrial ribosomal proteins in PyMT*^TrnE^* tumor cells. Bar graphs **(i)** showing quantification of Complex I-V, HSP60, TOMM40, TOMM20, MRPL28, and MRPS30 expression across PyMT*^TrnE^* Low, Mid, and High-mutant groups. β-Actin serves as a loading control. n = 4-5 biological replicates. **j, k** Immunoblots **(j)** showing expression levels of mitochondrial OXPHOS proteins, mitochondrial transport and folding proteins, and mitochondrial ribosomal proteins in PyMT*^ND5^* tumor cells. Bar graphs **(k)** showing quantification of Complex I-V, HSP60, TOMM40, TOMM20, MRPL28, and MRPS30 expression across PyMT*^ND5^* Low, Mid, and High-mutant groups. β-Actin serves as a loading control. n = 4-5 biological replicates. **l** Representative brightfield images (left) and size quantification (right) of tumor spheroids derived from PyMT*^WT^*, PyMT*^TrnE^*Mid, and PyMT*^ND5^* Mid tumor cells, with or without doxycycline treatment (PyMT*^WT^*, n = 163; PyMT*^WT^*+Dox, n = 173; PyMT*^TrnE^*Mid, n = 44; PyMT*^TrnE^* Mid+Dox, n = 65; PyMT*^ND5^* Mid, n = 115; PyMT*^ND5^* Mid+Dox, n = 142 tumoroids from 5 independent drug treatments) Data are as mean ± s.e.m. One-way ANOVA with Tukey’s multiple comparison test (**i, k, l**). Schematic in **g** was created in BioRender. NS, not significant; \**P* < 0.05; \*\**P* < 0.01; \*\*\**P* < 0.001; \*\*\*\**P* < 0.0001.

Based on these observations, we hypothesized that alterations in mitochondrial translation due to mtDNA mutations may represent a key determinant of tumor fate. We then analyzed the expression of proteins related to the large and small subunits of mitochondrial ribosomes in the proteomic dataset. Notably, MRPL and MRPS proteins were significantly upregulated in tumors from moderate-mutant groups, whereas no such increase was observed in tumors from high-mutant groups, suggesting enhanced mitochondrial translation in the moderate-mutant group (Fig. 4f, g). We further validated the expression levels of MRPL28 and MRPS30 in primary tumor cells, finding these proteins were highly expressed in moderate-mutant tumor cells compared to those with low and high mutation levels. Additionally, the expression of mitochondrial complexes I-V was upregulated in moderate-mutant tumor cells, particularly complexes II, III, and IV. Besides these, expression levels of mitochondrial transport- and folding-related proteins (TOMM20, TOMM40, HSP60) were also markedly elevated in moderate-mutant tumor cells (Fig. 4h-k). To confirm the functional significance of enhanced mitochondrial translation, we treated primary tumor cells *in vitro* with doxycycline, a compound that inhibits mitochondrial translation and affects protein synthesis. The doxycycline treatment of 3D cultured tumor spheres significantly reduced the size of moderate-mutant tumor spheres, indicating that suppression of mitochondrial translation substantially attenuates the growth advantage conferred by moderate mutations (Fig. 4l).

To determine whether the increased mitochondrial translation occurs prior to the tumor growth advantage in moderate-mutant groups, we dissected tumors at the initiation stage from PyMT-*TrnE* Low, Mid, and High mouse models at 20 weeks of age, when tumor weights showed similar sizes across groups (Fig. 5a). ScRNA-seq analysis of these primary tumors from different groups identified 14 distinct cell subpopulations based on marker gene expression, with the tumor cell subpopulation being the most abundant (Fig. 5b, c). DEG analysis of early-stage tumor cells showed that, compared to low- and high-mutant tumor cells, moderate-mutant tumor cells significantly upregulated genes encoding cytosolic ribosomal proteins (RPL and RPS), mitochondrial ribosomal proteins (MRPL and MRPS), and OXPHOS-related proteins (Fig. 5d). Pathway enrichment analysis further revealed significant activation of pathways related to ribosomal subunit assembly, ribosome biogenesis, mitochondrial respiratory chain function, and purine nucleotide metabolic process in moderate-mutant tumor cells (Fig. 5e). We refer to this phenomenon as the "ribosomal storm" in early-stage moderate-mutant tumor cells. Immunofluorescence staining confirmed elevated expression of RPL38 in moderate-mutant tumor cells (Fig. 5f, g), while immunoblots analysis showed increased levels of RPS27 in moderate-mutant tumor cells (Fig. 5h, i). Furthermore, early moderate-mutant tumor cells exhibited increased expression of Complex II, mitochondrial transport protein TOMM20, and mitochondrial ribosomal protein MRPL28 (Fig. 5h, i), findings consistent with those observed in late-stage tumor cells.

**Fig. 5.**
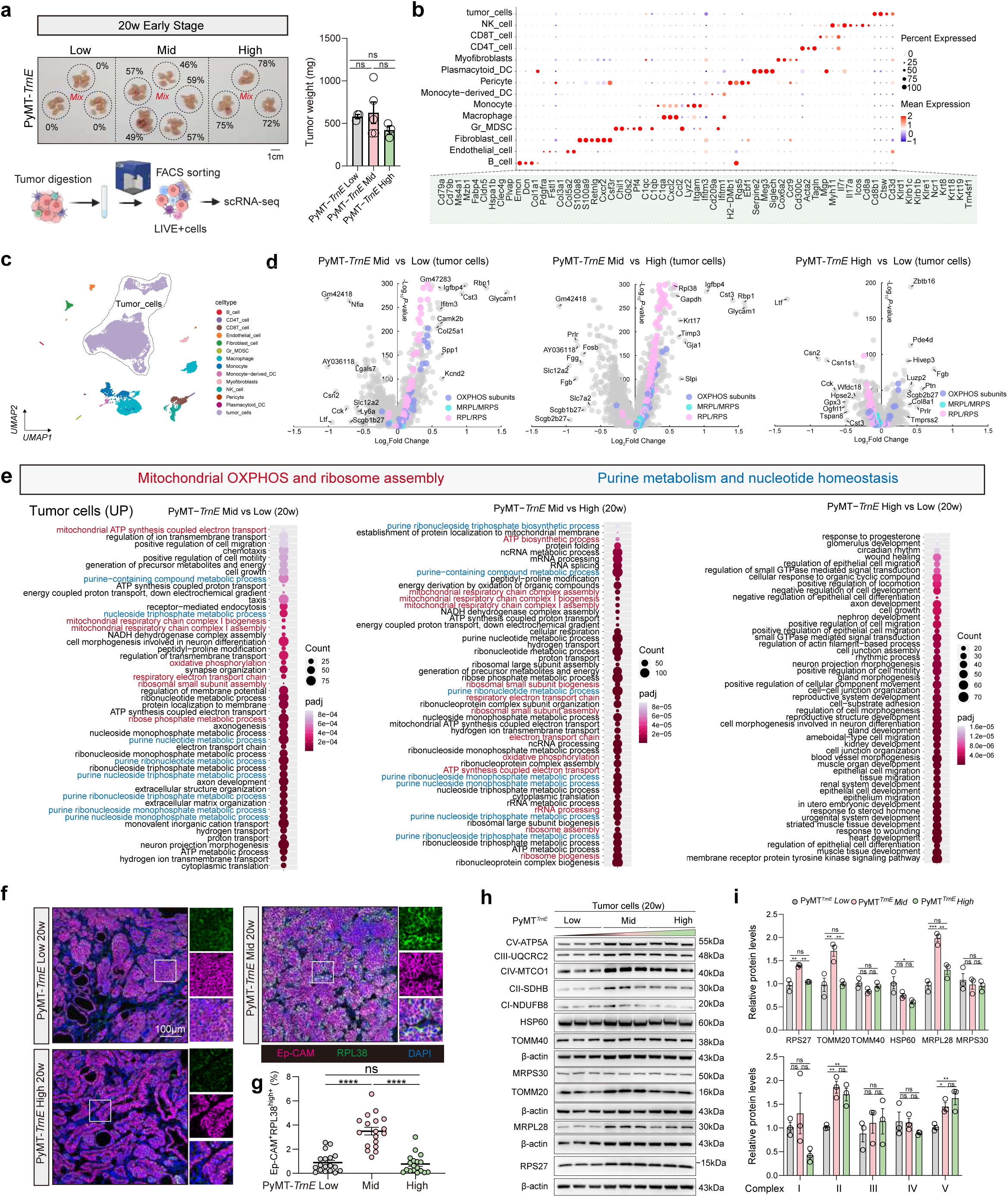
Moderate mtDNA heteroplasmy drives early activation of ribosome biogenesis in tumor cells. **a** Representative tumor images (top left), including PyMT-*TrnE* Low, Mid, and High-mutant groups (mtDNA mutation loads indicated on the graph), and tumor weight quantification (right) at 20 weeks. n = 3, 5, and 3 mice for PyMT-*TrnE* Low, Mid, and High groups, respectively. Schematic illustration (bottom left) showing the workflow for tumor dissociation and live cell sorting for scRNA-seq. **b** Gene expression from scRNA-seq experiments characterizing lineage-defining markers across cell clusters in PyMT-*TrnE* 20w tumor models. **c** Integrated Uniform Manifold Approximation and Projection (UMAP) plot and cell annotation of scRNA-seq data from tumors of PyMT-*TrnE* Low, Mid, and High groups at 20 weeks. **d** Volcano plots of tumor cell clusters from scRNA-seq data at 20 weeks showing top differentially expressed genes between groups, with OXPHOS subunits labeled in purple, MRPL/MRPS in blue, and RPL/RPS in pink. Comparisons include PyMT-*TrnE* Mid vs Low (left), PyMT-*TrnE* Mid vs High (center), and PyMT-*TrnE* High vs Low (right). Genes with FDR-adjusted two-tailed t-test *P* < 0.05 were considered statistically significant. **e** Bubble plot showing pathway enrichment in tumor cell clusters from 20w PyMT-*TrnE* Low, Mid, and High tumors, with comparisons of Mid vs Low (top left), Mid vs High (top right), and High vs Low (bottom), where bubble size represents gene count and color intensity indicates adjusted p-value (padj). **f, g** Representative immunofluorescence staining **(f)** and quantification **(g)** of high RPL28 expression in tumor cells (Ep-CAM^+^) from PyMT-*TrnE* Low, Mid, and High mice at 20 weeks of age (n = 18 fields from 3 mice per group). Scale bars, 100 μm. **h, i** Immunoblots **(h)** showing expression levels of ribosomal protein, mitochondrial OXPHOS proteins, mitochondrial transport and folding proteins, and mitochondrial ribosomal proteins in PyMT*^TrnE^* tumor cells at the early 20-week stage. Bar graphs **(i)** showing quantification of RPS27, Complex I-V, HSP60, TOMM40, TOMM20, MRPL28, and MRPS30 expression across PyMT*^TrnE^* Low, Mid, and High-mutant groups at 20 weeks. β-Actin serves as a loading control. n = 3 biological replicates. Data are as mean ± s.e.m. One-way ANOVA with Tukey’s multiple comparison test (**a, g, i**). Schematic in **a** (bottom left) were created in BioRender. NS, not significant; \**P* < 0.05; \*\**P* < 0.01; \*\*\**P* < 0.001; \*\*\*\**P* < 0.0001.

Collectively, these results demonstrate that moderate-mutant tumor cells gain a growth advantage through enhanced mitochondrial protein expression driven by the “ribosomal storm” and increased mitochondrial translation.

### LARS2 upregulation defines a moderate-mutant tumor cell–intrinsic dependency on mitochondrial translation

Given the evidence of enhanced mitochondrial translation in moderate-mutant tumor cells, we next sought to identify upstream regulators of this process. Single cells were isolated from tumor tissues of multiple PyMT-*TrnE*/*ND5* mouse groups for scRNA-seq analysis, which identified 16 distinct cell populations (Fig. 6a, b, and Extended Data Fig. 5a). We compared the gene expression profiles from the scRNA-seq of tumor cells from the PyMT-*TrnE*/*ND5* Mid and High-mutant groups. A combination of volcano plots and Venn diagram analyses revealed 3,420 overlapping DEGs, encompassing both upregulated and downregulated transcripts. Among these, *Lars2* emerged as significantly upregulated in tumor cells in the moderate-mutant groups (Fig. 6c-e). The *Lars2* gene encodes leucyl-tRNA synthetase 2, a mitochondrial enzyme essential for mitochondrial protein synthesis ^21^ (Fig. 6f). In our scRNA-seq analysis, we detected 17 cytosolic and 17 mitochondrial aminoacyl-tRNA synthetases in tumor cells. Notably, only *Lars2* exhibited significant upregulation in moderate-mutant cells, while the expression levels of all other synthetases, both cytosolic and mitochondrial, remained largely unchanged (Extended Data Fig. 4a). Flow cytometric analysis corroborated this finding, demonstrating a significant increase in the frequency of LARS2-high cells among moderate-mutant tumor cells (Fig. 6g). Notably, no significant difference in mutation load was observed between LARS2-high and LARS2-low subpopulations (Extended Data Fig. 4b).

**Fig. 6.**
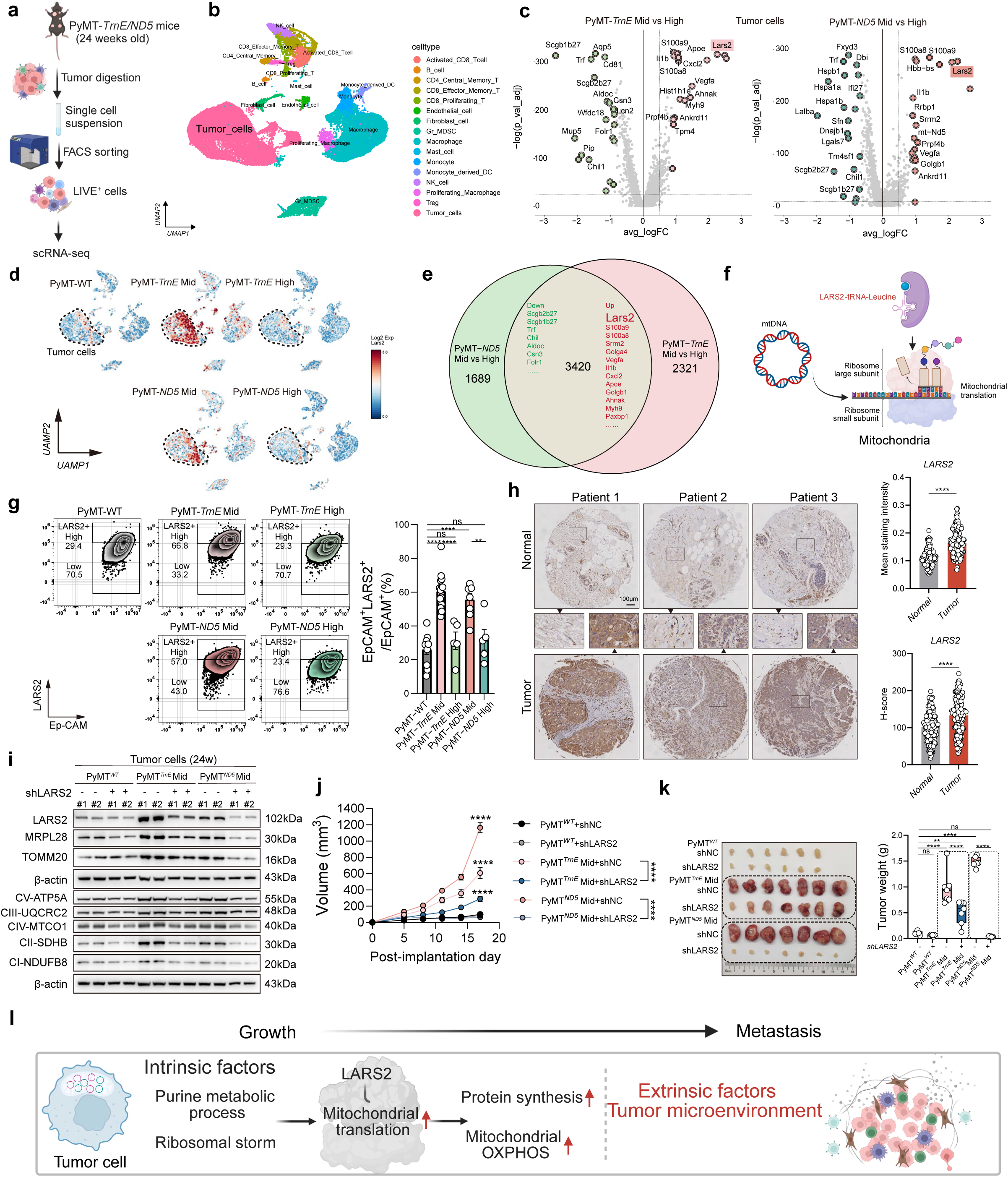
LARS2-dependent mitochondrial translation defines a therapeutic vulnerability in tumors with moderate mtDNA heteroplasmy. **a** Schematic of single-cell RNA sequencing from tumors of PyMT-*TrnE*/*ND5* mice. The live cells were sorted by flow cytometry and combined in equal numbers within the same group for 10X genomic scRNA-seq (n = 1 pooled from three mice per group). **b** Integrated UMAP plot and cell annotation of scRNA-seq data from tumors of PyMT-WT, PyMT-*TrnE*, and PyMT-*ND5* groups. **c** Volcano plots of tumor cell clusters from scRNA-seq data at 24 weeks showing differentially expressed genes between groups (Green: down, red: up). Comparisons include PyMT-*TrnE* Mid vs High (left) and PyMT-*ND5* Mid vs High (right). Differences were considered statistically significant at FDR-adjusted *P* < 0.05 using two-tailed t-tests. **d** UMAP visualization based on scRNA-seq data showing cluster-specific expression of *Lars2* in PyMT-WT, PyMT-*TrnE* Mid/High, and PyMT-*ND5* Mid/High groups. **e** Venn diagrams depicting the overlap and unique sets of differentially expressed genes in tumor cell clusters from 24-week scRNA-seq, comparing PyMT-*TrnE* Mid vs High and PyMT-*ND5* Mid vs High groups. **f** Schematic showing the role of LARS2 in mtDNA-mediated translation. LARS2 is a mitochondrial leucyl-tRNA synthetase that catalyzes the charging of mitochondrial tRNA^Leu (UUR) with leucine, a critical step in mitochondrial protein synthesis. **g** Representative flow cytometry plots (left) and bar plots (right) showing the percentages of Ep-CAM^+^ LARS2^+^ tumor cells of PyMT-WT, PyMT-*TrnE* Mid/High PyMT- *ND5* Mid/High group. n = 10, 15, 5, 8 and 5 mice for PyMT-WT, PyMT-*TrnE* Mid/ High, PyMT- *ND5* Mid/ High group, respectively. **h** Representative immunohistochemical staining of LARS2 in breast cancer tissue microarrays (Normal, n = 164; Tumor, n = 164) (left), and corresponding quantification (mean intensity, top right; H-score, bottom right). **i** Immunoblots showing expression levels of mitochondrial OXPHOS proteins (Complex I-V), mitochondrial transport proteins (TOMM20), and mitochondrial ribosomal proteins (MRPL28) in PyMT*^WT^*, PyMT*^TrnE^*Mid, and PyMT*^ND5^* Mid tumor cells with or without *Lars2* knockdown. β-Actin serves as a loading control. n = 4 biological replicates. **j, k** Tumor growth curves **(j)** following implantation of *Lars2* non-knockdown and knockdown tumor cells from PyMT*^WT^*, PyMT*^TrnE^*Mid, and PyMT*^ND5^* Mid groups. Tumor images (left) and tumor weight quantification (right) after dissection **(k)** (n = 7 mice per group). **l** Conceptual model linking tumor-intrinsic mitochondrial reprogramming to extrinsic immune evasion, culminating in the progression from tumor growth to metastasis. Data are as mean ± s.e.m. two-way ANOVA (**j**) and One-way ANOVA with Tukey’s multiple comparison test (**g, k**). Student’s two-sided unpaired t-test (**h**). Schematic in **a, f** and **l** were created in BioRender. NS, not significant; \**P* < 0.05; \*\**P* < 0.01; \*\*\**P* < 0.001; \*\*\*\**P* < 0.0001.

We further assessed *Lars2* expression across normal tissues, primary tumor tissues, and metastatic lesions using human breast cancer datasets from the GEO, GTEx, TCGA, and TARGET databases. Our analysis revealed that *Lars2* was markedly overexpressed in primary tumors compared to normal tissues (Extended Data Fig. 4c). Consistent with this, immunohistochemical staining of primary breast cancers and adjacent normal tissues showed significantly elevated LARS2 signals in tumor tissues, as quantified by both mean staining intensity and H-score (Fig. 6h). Survival analysis based on the PAM50 molecular subtypes of breast cancer (Normal, Luminal A/B, HER2-enriched, Basal-like) revealed that high *Lars2* expression was associated with significantly reduced overall survival specifically in patients with the Basal-like subtype, the most aggressive form of breast cancer, whereas no significant effect was observed in the Normal-like subtype (Extended Data Fig. 4d). Expanding our analysis to additional cancer types, we found that *Lars2* was not only overexpressed in breast cancer, but also significantly upregulated in most other tumor types examined (Extended Data Fig. 4e). To evaluate the functional consequences of *Lars2* upregulation in moderate-mutant tumor cells, we established *Lars2* knockdown cell lines using primary tumor cells derived from PyMT-WT/*TrnE*/*ND5* models *in vitro*. *Lars2* inhibition effectively attenuated the elevated expression of mitochondrial respiratory complexes I-V induced by moderate mutations, with the most pronounced reduction observed in complex II levels. Additionally, knockdown of *Lars2* significantly diminished the expression of the mitochondrial import receptor TOMM20 and the mitochondrial ribosomal protein MRPL28 (Fig. 6i). *In vivo* xenograft experiments further demonstrated that *Lars2* knockdown markedly suppressed tumor growth driven by moderate mutations (Fig. 6j, k). Collectively, our findings establish *Lars2* as a key regulator in moderate-mutant tumor cells that governs mitochondrial translation, thereby modulating tumor progression. Given the significant challenges associated with manipulating mtDNA mutation levels *in vivo*, *Lars2* represents a promising alternative therapeutic target for future cancer treatments.

### Moderate mtDNA mutations remodel the tumor immune microenvironment

The enhanced proliferative capacity of moderate-mutant tumor cells is accompanied by a striking increase in metastasis—suggesting that intrinsic metabolic advantages alone are insufficient to explain their aggressive phenotype. We therefore asked how these cells remodel the tumor microenvironment to enable immune evasion and systemic dissemination (Fig. 6l). To address this, we analyzed the immune landscape across mtDNA mutation burdens.

Interestingly, an immunosuppressive condition was already evident in primary tumors of the moderate-mutant group at an early stage—when tumor size did not differ from controls — characterized by an increased proportion of myeloid-derived suppressor cells (MDSCs) ^22^ and a decreased proportion of CD8⁺ T cells compared to low- and high-mutant groups (Fig. 7a). Myeloid cells in these tumors displayed enhanced expression of anti-inflammatory genes and upregulated pathways associated with neutrophil migration (Fig. 7b), and CD8⁺ T cells exhibited significantly lower activation and migration scores, consistent with reduced infiltration (Fig. 7b).

**Fig. 7.**
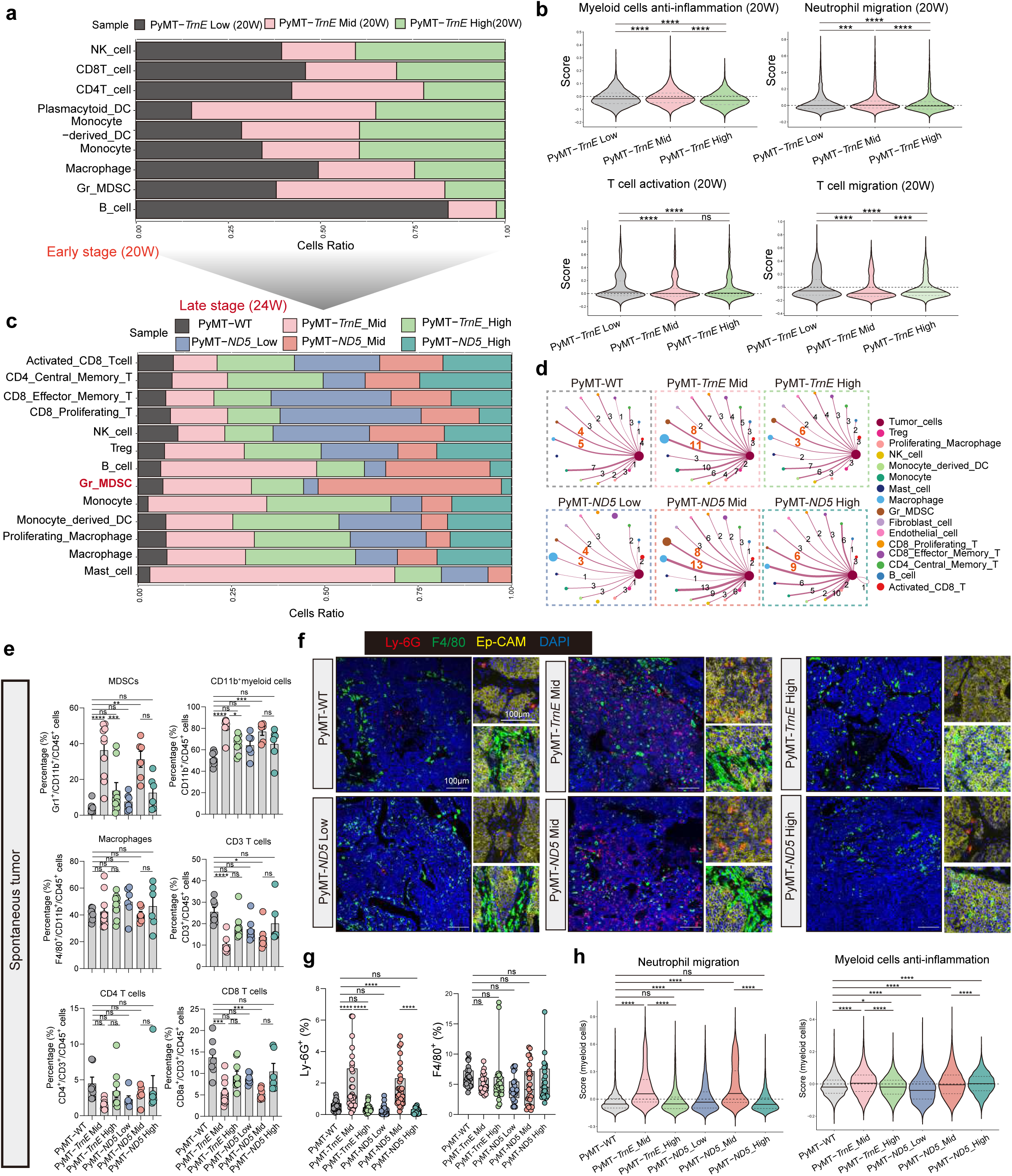
mtDNA mutations drive progressive establishment of an immunosuppressive microenvironment from early to late tumorigenesis. **a** Proportions of immune subpopulations in tumors of PyMT-*TrnE* Low/Mid/ High group at 20w. **b** Transcripts from myeloid cells in tumors were assessed for enrichment of gene signatures for neutrophil migration and anti-inflammation. Transcripts from T cells in tumors were assessed for enrichment of signatures of T cells activation and migration. **c** Proportions of immune subpopulations in PyMT-WT, PyMT-*TrnE* Mid/ High, PyMT-*ND5* Low/Mid/ High group at 24w. **d** Analysis of tumor cells, MDSCs, and macrophages in PyMT-WT, PyMT-*TrnE* Mid/High, and PyMT-*ND5* Low/Mid/High tumors showing distinct types of receptor–ligand interactions between these cell populations. **e** Bar graphs showing the percentage of indicated immune cell types across groups. n = 7, 10, 9, 6, 6 and 6 mice for PyMT-WT, PyMT-*TrnE* Mid/High, PyMT- *ND5* Low/Mid/High group, respectively. **f, g** Representative immunofluorescence staining **(f)** and quantification **(g)** of MDSCs and macrophages in tumors from PyMT-WT, PyMT-*TrnE* Mid/High, PyMT-*ND5* Low/Mid/ High mice (MDSCs: n = 30 fields from 6 mice per group; macrophages: n = 26, 27, 26, 24, 27, 24 fields from 6 mice, respectively). Scale bars, 100 μm. **h** Transcripts from myeloid cells in tumors were assessed for enrichment of gene signatures for neutrophil migration and anti-inflammation. Data are as mean ± s.e.m. Kruskal-Wallis test with Dunn’s multiple comparisons test (**b, h**) and One-way ANOVA with Tukey’s multiple comparison test (**e, g**). DC, dendritic cell. NK cell, Natural Killer Cell. NS, not significant; \**P* < 0.05; \*\**P* < 0.01; \*\*\**P* < 0.001; \*\*\*\**P* < 0.0001.

In late-stage tumors, this immunosuppressive landscape persisted, with moderate-mutant tumors again showing elevated MDSCs and diminished CD8⁺ T cell infiltration compared to both PyMT-WT and high-mutant counterparts (Fig. 7c). To investigate how tumor cells communicate with the immune compartment, we analyzed receptor–ligand interactions, revealing enhanced crosstalk between moderate-mutant tumor cells and MDSCs as well as macrophages (Fig. 7d). These observations were validated by flow cytometry, which revealed elevated infiltration of MDSCs and myeloid cells, along with decreased CD8^+^ T cell infiltration, in the moderate-mutant group (Fig. 7e). Immunofluorescence staining confirmed higher MDSC levels in moderate-mutant tumors, while macrophage proportions remained comparable across groups (Fig. 7f, g). Transcriptomic analysis of myeloid cells showed significant enrichment of the neutrophil migration gene signature in the moderate-mutant group (Fig. 7h), consistent with increased MDSCs infiltration. Additionally, these cells displayed an enhanced anti-inflammatory gene signature, as indicated by significantly higher enrichment scores for anti-inflammatory genes (Fig. 7h).

Spleen size was markedly increased in the moderate-mutant group (Extended Data Fig. 5b), and similar immune cell distribution patterns were observed in the spleen, with elevated MDSC levels and reduced CD8^+^ T cell proportions (Extended Data Fig. 5c). However, no significant differences in peripheral blood immune cell composition were detected among the groups (Extended Data Fig. 5d).

Collectively, these results indicate that moderate mtDNA mutations are associated with the development of an immunosuppressive tumor microenvironment. Moreover, immunosuppressive signaling in the tumor microenvironment is initiated at an early stage, coinciding with these intrinsic translational adaptations.

### Moderate-mutant tumor cells establish an immunosuppressive tumor microenvironment through upregulation of S100A8/A9 signaling

To determine whether the immunosuppressive microenvironment is driven by tumor cell–intrinsic factors, we analyzed the immune profiling data from tumors formed by transplanted moderate-mutant primary tumor cells or mutant 4T1 cells, which showed a significant increase in MDSCs and a concomitant decrease in CD8^+^ T cells (Extended Data Fig. 6a, b), recapitulating the immune landscape observed in PyMT-*TrnE*/*ND5* spontaneous tumors. Second, *in vitro* co-culture assays demonstrated that moderate-mutant primary tumor cells exhibited enhanced neutrophil recruitment, whereas high-mutant cells showed recruitment levels similar to or even lower than controls (Extended Data Fig. 6c). Third, bulk RNA-seq analysis of transplanted mutant 4T1-derived tumors further revealed significant upregulation of genes associated with neutrophil migration and extravasation (Extended Data Fig. 6d), collectively indicating a heightened capacity of moderate-mutant tumor cells to recruit neutrophils into the tumor microenvironment.

To identify upstream drivers, we reanalyzed scRNA-seq data from PyMT-*TrnE/ND5* tumors and found that *S100A8* and *S100A9*—key alarmins involved in myeloid cell recruitment ^23,24^—were among the top three upregulated genes, along with *Lars2* in tumor cells of the moderate-mutant group (Fig. 6e). Notably, we also observed increased expression of *Cxcl2* transcript level, a chemokine implicated in MDSC trafficking ^25^ (Extended Data Fig. 6e), further supporting the enhanced recruitment of MDSCs by tumor cells. The analysis of mtscATAC-seq data showed increased chromatin accessibility at the *S100A9* locus in PyMT-*TrnE*/*ND5* Mid tumors (Extended Data Fig. 6f), which correlated with elevated *S100A9* mRNA and serum protein levels (Extended Data Fig. 6g, h). Upregulation of S100A8/A9 was also confirmed in differentially expressed genes (DEGs) from tumors derived from transplanted moderate-mutant 4T1 cells (Extended Data Fig. 6i, j). Immunofluorescence staining revealed abundant S100A9 signals in tumors originating from both transplanted moderate-mutant 4T1 and primary tumor cells (Extended Data Fig. 6k, l).

Given the central role of S100A8/A9 in shaping immunosuppressive niches, we tested whether targeting this pathway could disrupt tumor progression (Fig. 8a). Administration of the S100A9 inhibitor tasquinimod significantly suppressed tumor growth and metastasis in models driven by moderate-mutant cells, but had minimal effect on high-mutant tumors (Fig. 8b-e). Tasquinimod also reduced MDSCs infiltration in moderate-mutant tumors (Fig. 8f, g), supporting its functional relevance. To assess the contribution of MDSCs to tumor progression, we performed *in vivo* depletion using anti-Gr-1 antibodies (Fig. 8h). While MDSC depletion effectively reduced their abundance without altering T cell proportions (Fig. 8i), primary tumor growth was unaffected—though a modest reduction was observed in transplanted 4T1-*TrnE* Mid tumors (Fig. 8h, j). Notably, MDSCs depletion led to a significant decrease in metastasis in the 4T1-*TrnE*/*ND5* Mid group (Fig. 8k). Critically, *Lars2* knockdown was associated with a marked decrease in the proportion of MDSCs within the tumor microenvironment (Extended Data Fig. 7a), and reduced S100A9 signals were consistently observed in *Lars2*-knockdown tumors, coinciding with attenuated MDSC accumulation (Extended Data Fig. 7b, c).

**Fig. 8.**
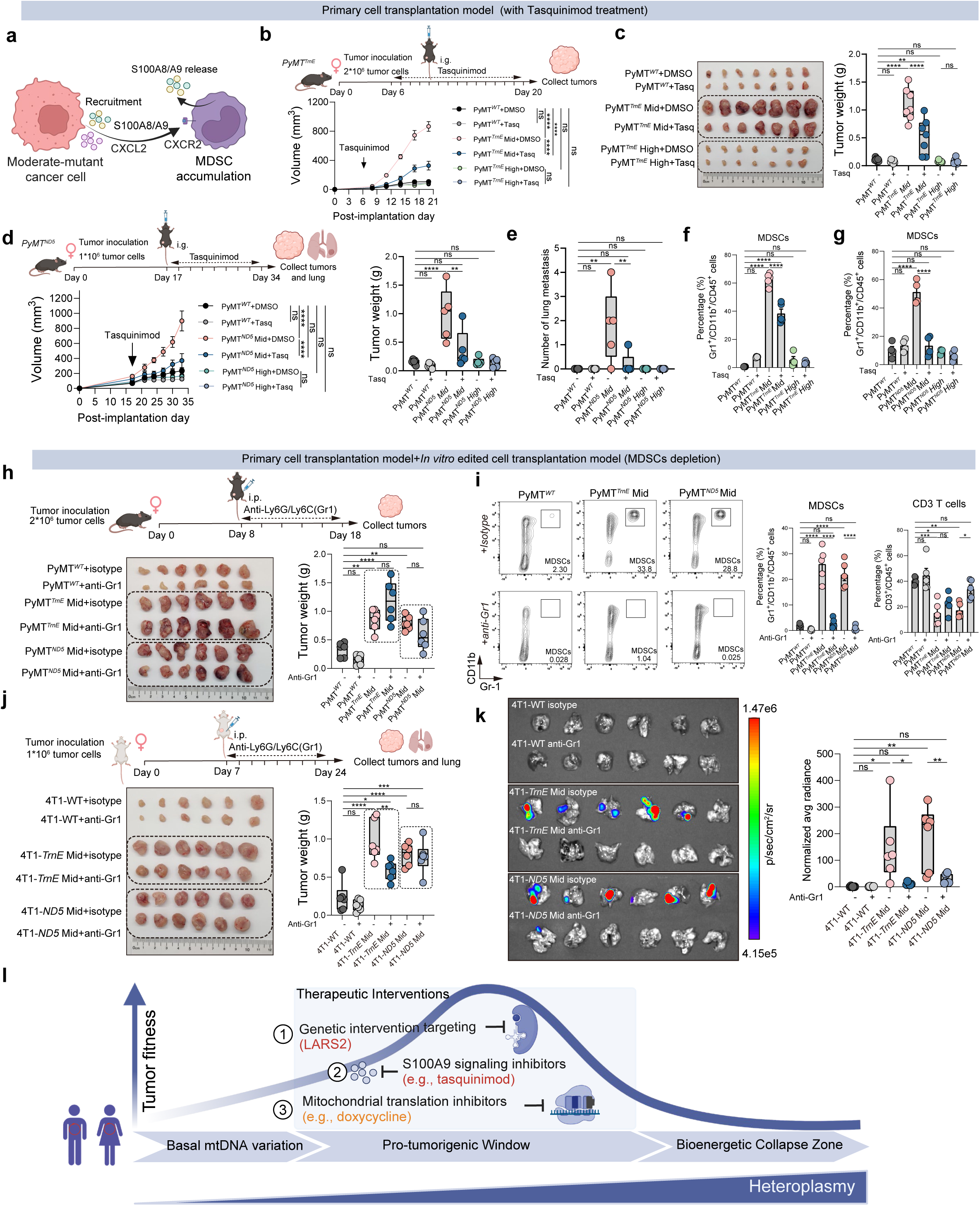
Targeting the S100A8/A9–MDSCs axis suppresses tumor progression in moderate-mutant models. **a** Schematic of the secretion and crosstalk of S100A8/A9 and CXCL2 signaling between tumor cells and MDSCs in the tumor microenvironment. **b, c** PyMT*^WT^*, PyMT*^TrnE^* Mid/High cells were inoculated subcutaneously into wild-type mice, followed by treatment with tasquinimod or DMSO control every day from day 6 post-inoculation, Growth curves are shown **(b)**. Images of dissected tumors (**c,** left) and corresponding tumor weight quantification (**c,** right) (n = 7 mice per group). **d, e** PyMT*^WT^*, PyMT*^ND5^* Mid/High cells were inoculated subcutaneously into wild-type mice, followed by treatment with tasquinimod or DMSO control every day from day 17 post-inoculation, Growth curves **(d**, left), tumor weight (**d**, right) and lung metastasis quantification **(e)** (n = 5 mice per group) are shown. **f, g** Tumors were harvested for flow cytometric analysis, showing the proportion of MDSCs in PyMT*^TrnE^* Mid/High **(f)** or PyMT*^ND5^* Mid/High **(g)** treatment group. (n = 4-5 mice per group) **h** PyMT*^WT^*, PyMT*^TrnE^* Mid and PyMT*^ND5^* Mid cells were inoculated subcutaneously into wild-type mice, followed by treatment with anti-Ly6G/Ly6C (Gr1) or isotype control every other day from day 8 post-inoculation. Images of dissected tumors (left) and corresponding tumor weight quantification (right) (n = 6 mice per group). **i** Representative flow cytometry plots (left) showing the proportion of MDSCs and the percentage of indicated immune cell populations (right) in tumors derived from PyMT*^WT^*, PyMT*^TrnE^* Mid and PyMT*^ND5^*Mid cells following anti-Ly6G/Ly6C (Gr1) treatment, as determined by flow cytometry (n = 5 mice per group). **j** 4T1-WT, 4T1-*TrnE* Mid, 4T1-*ND5* Mid cells were inoculated subcutaneously into wild-type mice, followed by treatment with anti-Ly6G/Ly6C(Gr1) or isotype control every other day from day 7 post-inoculation. Images of dissected tumors (left) and corresponding tumor weight quantification (right) (n = 6 mice per group). **k** Bioluminescence imaging (left) and quantification (right) showing lung metastasis in wild-type mice inoculated with luciferase-labeled 4T1-WT, 4T1-*TrnE* Mid, and 4T1-*ND5* Mid cells, followed by treatment with anti-Ly6G/Ly6C (Gr-1) or isotype control every other day starting from day 7 post-inoculation (n = 6 mice per group). **l** Graphical model depicting the impact of mtDNA heteroplasmy on tumor fate. The level of mutant mtDNA determines tumor progression through a threshold-dependent mechanism. Basal mtDNA variation (low heteroplasmy, 0–30%) exerts minimal impact on mitochondrial function and tumor behavior. Moderate heteroplasmy (30–60%) defines a pro-tumorigenic window characterized by LARS2-driven mitochondrial translational reprogramming, S100A8/A9 secretion, and MDSCs-mediated immunosuppression. This state creates a therapeutic vulnerability that can be effectively targeted by *Lars2* knockdown, inhibition of mitochondrial translation, or blockade of S100A9 signaling. In contrast, high heteroplasmy (>70%) leads to severe OXPHOS deficiency and bioenergetic collapse, impairing tumor growth despite strong oncogenic drivers. Data are as mean ± s.e.m. two-way ANOVA (**b, d**) and One-way ANOVA with Tukey’s multiple comparison test (**c, d, e, f, g, h, i, j, k**). Schematic in **a, b, d, h, j, l** were created in BioRender. NS, not significant; \**P* < 0.05; \*\**P* < 0.01; \*\*\**P* < 0.001; \*\*\*\**P* < 0.0001.

These findings unveil a unified mechanism through which mtDNA heteroplasmy modulates tumor fate in a mutation load-dependent manner, orchestrating a coordinated program of intrinsic metabolic adaptation and extrinsic immunosuppressive niche formation. We propose a model in which moderate levels of mutant mtDNA—sufficiently high to alter cellular physiology but not to induce bioenergetic failure—establish a pro-tumorigenic window. This window can be effectively suppressed by either *Lars2* knockdown, inhibition of mitochondrial translation or inhibition of S100A9. In contrast, tumors harboring high mutant levels experience bioenergetic collapse, thereby impairing their capacity to sustain tumor progression. Collectively, our study reveals that tumor cells actively sense mitochondrial genome dosage to coordinate metabolic fitness with immune evasion. This paradigm positions mtDNA heteroplasmy not merely as a passive biomarker of dysfunction, but as a programmable regulator of tumor behavior, with broad implications for targeting the LARS2 –mitochondrial translation–S100A8/A9 axis in cancer therapy (Fig. 8l).

## Discussion

By establishing two novel mouse models (PyMT-*TrnE/ND5*), we provide direct *in vivo* evidence that moderate levels of mtDNA mutations—previously considered to have limited clinical significance ^26–29^—are active modulators of tumor growth and metastasis, whereas high mutant loads suppress tumorigenesis. Our findings showed that in the context of cancer, these otherwise “silent” mutations can be reactivated, enhancing tumor cell fitness. This reframes mtDNA heteroplasmy not as a passive biomarker of genomic instability, but as a tunable regulator of tumor fate—one that integrates bioenergetic state with cell-extrinsic signalling to license immune evasion and metastasis.

Furthermore, we defined a "pro-tumorigenic window" in which moderate-mutant level of mtDNA mutations confer a selective advantage to cancer cells. This apparent “self-rescuing” mechanism in moderate-mutant tumor cells paradoxically contributes to its progression. In contrast, high mtDNA mutation burdens are associated with impaired respiratory chain function, reduced bioenergetic capacity, and restricted tumor growth—findings that align with clinical observations showing no increased cancer incidence among individuals with mitochondrial disorders ^30^. These insights have raised concerns regarding therapeutic strategies aimed at reducing mtDNA mutation load ^31–33^: although such approaches may benefit patients with mitochondrial diseases, they could simultaneously eliminate a natural “protective barrier” in pre-neoplastic tissues, thereby unmasking latent pro-tumorigenic potential.

Our data further highlight the dynamic nature of mtDNA heteroplasmy during tumor evolution. While moderate-mutant mtDNA mutations appear to confer early adaptive advantages, the nonsynonymous mutation in the *ND5* gene is progressively eliminated as tumors expand, leading to a decline in mutant levels. This pattern indicates strong purifying selection during tumor progression—initially permissive of moderate-mutant level but ultimately favoring metabolic efficiency in advanced stages. Consequently, the low level of the nonsynonymous mutations observed in late-stage tumors likely reflects a later adaptive shift rather than representing the initial pro-tumorigenic event. This temporal evolution underscores the need for a re-evaluation of how mtDNA mutations are interpreted in clinical genomic analyses.

Mechanistically, we identified a defining characteristic of tumors harboring moderate mtDNA mutations: the early and sustained activation of ribosomal pathways, which we designate as a “ribosomal storm” and propose as a compensatory adaptive response. Consistent with recent findings of recurrent, cancer-associated mutations in mitochondrial rRNA genes (MT-RNR1/2) ^34^, our results position mitochondrial translation as a central node in mtDNA-driven tumorigenesis. Central to this phenomenon is LARS2, the mitochondrial leucyl-tRNA synthetase and a key regulator of mitochondrial protein synthesis ^21,35,36^. Given that leucine is among the most frequently encoded amino acids in mtDNA ^37,38^, LARS2 serves as a critical determinant of translational fidelity within the mitochondria. This functional dependency establishes a therapeutic vulnerability—tumors exhibiting reliance on this pathway demonstrate heightened sensitivity to LARS2 inhibition—thereby revealing a potential precision therapeutic strategy for cancers driven by mtDNA mutations. Moreover, moderate mtDNA mutations reshape the tumor microenvironment. They suppress antigen presentation and elevate expression of S100A8/A9, leading to recruitment of MDSCs and consequent immunosuppression ^39,40^. These cells have been proved to facilitate immune evasion, vascular occlusion, and nutrient restriction ^41–43^, thereby establishing a permissive ecological niche for tumor progression. Collectively, these findings open new avenues for immunotherapeutic interventions targeting the S100A9 pathway or MDSCs.

In summary, our findings demonstrate that mtDNA mutations drive a multifaceted adaptive program encompassing bioenergetic resilience, enhanced translational capacity, and immune remodelling (Extended Data Fig. 7d). This adaptive response establishes mtDNA heteroplasmy as a non-nuclear determinant of tumor progression, underscoring the critical importance of considering both tumor phase and heteroplasmy level when evaluating mtDNA variants in clinical tumor samples.

## Methods

### Animal models

All procedures followed university guidelines and were approved by the Animal Use and Care Committee of Westlake University (Approval No. 22-003-5-JM-3). Mice were maintained in a specific pathogen-free (SPF) facility at Westlake University under controlled conditions (temperature: 22 ± 1 °C; relative humidity: 40-70%) with a 12-hour light/dark cycle. Standard chow and water were provided ad libitum. All mice were backcrossed to the C57BL/6 background. Six- to eight-week-old female C57BL/6 mice were purchased from the Animal Use and Care Committee of Westlake University. Six- to eight-week-old female BALB/cJ mice were purchased from Beijing Vital River Laboratory Animal Technology Co., Ltd. *MMTV*-PyMT transgenic mice (FVB/N-Tg(*MMTV*-PyVT)634Mul/J, JAX002374) were backcrossed to C57BL/6J mice for more than 10 generations. *TrnE* (mtDNA G14102A single mutant model) /*ND5* (mtDNA G12918A single mutant model) point mutant mice were constructed previously ^17,18^. Male *MMTV*-PyMT mice were crossed with female *TrnE* and *ND5* mutant mice to generate PyMT-*TrnE* and PyMT-*ND5* mice. The PyMT allele can be distinguished from the wild-type allele by PCR using the primer sets: PyMT_Forward: 5′- GGAAGCAAGTACTTCACAAGGG -3′, PyMT_Reverse: 5′- GGAAAGTCACTAGGAGCAGGG -3′.

Spontaneous tumor-bearing mice were analyzed for tumor size and lung metastasis at 20 and 24 weeks. Tumor volume was calculated using the formula (L × W)^2^/2, where L is length and W is width. Total tumor burden was calculated by summing individual tumor volumes of each mouse with an end-point defined when total burden reached 4,000 mm^3^ or one tumor reached 2,000 mm^3^. During tumor measurement, the investigators were blinded to the genotypes of mice.

### Mouse tumor dissection and primary cell culture

Mouse breast tumors were dissected and digested according to previous protocol with minor revisions ^44^. Mouse breast tumors were dissected into 9.5mL DMEM/F12 media (C11330500CP, Gibco), the tumor tissues were chopped with razor blade into 1mm^3^ small pieces, followed by digestion with collagenase type IV (1mg/mL) (#17104-019, Gibco), DNaseI (0.1mg/mL) (#LS002139, Worthington Biochemical) and hyaluronidase (0.2mg/mL) (#LS002592, Worthington Biochemical) at 37°C for 1 h. The digested tissue was filtered through a 70 μm cell strainer and then centrifuged in a 15 ml centrifuge tube at 1500rpm for 5 minutes. Red blood cells were lysed with 5 mL ACK (#00-4300-54, Invitrogen) at room temperature for 2 min, washed once with PBS. Dissociated cells were counted and resuspended in designated medium for subsequent staining and FACS analysis.

The primary cell pellets were resuspended in 10ml culture media with DMEM/F-12, GlutaMAX (#10565018, Gibco) + 10%FBS (#10091148, Gibco) + P/S antibiotics (Gibco) supplemented with EGF (10ng/uL, BD Bioscience), Y-27632 dihydrochloride (10uM, Tocris), Normocin (ant-nr-1, InvivoGen). The cell suspension was plated in a 10 cm^2^ dish and cultured in a humidified incubator at 37°C with 5% CO_2_. After overnight adherence, the cells were washed with PBS to remove non-adherent cells. To obtain a purer population of tumor cells, the adherent cells were trypsinized and the primary cancer cells were purified by flow cytometry using a cell-surface antibody (EpCAM, BioLegend).

### mtDNA mutant tumor cell lines

4T1 (mouse breast cancer cell line) (ATCC, CRL-2539) were obtained from Westlake University Cell Bank. Cells were cultured in RPMI 1640 (BasalMedia) supplemented with 10% FBS, 100 IU/ ml penicillin/streptomycin (Invitrogen). Cell cultures were frequently monitored for mycoplasma contamination, and only mycoplasma-negative cells were used for experiments.

The DdCBE plasmids were tested for efficient editing as described previously ^17,18^. To introduce single-site mtDNA mutations (*TrnE* G14702A mutation, *ND5* G12918A mutation) in 4T1 cells *in vitro*, 2.5 µg of each DdCBE plasmid pair were transfected into 6*10^5^ cells using the Lipofectamine 3000 Transfection Reagent (Invitrogen). Following transfection, cells were incubated for 72 hours before FACS was performed using a BD FACSAria Fusion Cell Sorter to isolate eGFP and mCherry double-positive cells, plated as single cells, and allowed to expand into colonies over a one-month period without further transfection or editing. These colonies were established as stable cell lines and employed for downstream analyses, including mutation load validation, and biochemical assays.

Luciferase-expressing lentivirus were purchased from GeneChem (Ubi-MCS-firefly_Luciferase-IRES-Puromycin). 4T1 cells (mutant or non-mutant) constructed above were transfected with lentivirus for 72 h and then screened with puromycin at a concentration of 1.5 µg/ ml to construct a cell line stably expressing luciferase.

### mtDNA mutation sequencing

Mice tissues (tail, blood, tumor, lung, liver) and cells were lysed with a lysis buffer containing 25mM NaOH and 0.2mM EDTA. Mutation load in mtDNA was quantified using Sanger or next-generation sequencing (NGS) based Hi-TOM sequencing ^45,46^. The accuracy of Hi-TOM has been previously validated ^17,18^. Mutations were detected using HiTOM analysis (http://www.hi-tom.net/hi-tom/). The primers used for mutation load measurement were as follows:

mouse *TrnE*-F: 5′- ggagtgagtacggtgtgcctaacccaagacaaccaacc -3′;

mouse *TrnE*-R: 5′- gagttggatgctggatggggttaataattttaaataatgggtgtg -3′;

mouse *ND5*-F: 5′- ggagtgagtacggtgtgcaatcataccattcacatcatc -3′;

mouse *ND5*-R:5′- gagttggatgctggatggacatagctgttatagaagtgg -3′;

Specifically, we performed mutation load screening on primary tumor cells isolated from mice with different mtDNA mutations. The tumors were digested and plated in separate dishes for culture. After the cells adhered, they were washed, trypsinized, and purified using flow cytometry with cell-surface antibodies (EpCAM, BioLegend). The mtDNA mutation load was then determined, and cells with the desired mutation loads were selected, expanded in culture, and used for subsequent experiments.

### The generation of knockdown cell lines

Stable knockdown primary cancer cells were generated via lentiviral transduction. The lentiviral vectors encoding shRNAs against *Lars2* and the nonspecific control were purchased from Tsingke Biotech. Cells were selected using 2 µg/ ml puromycin. The knockdown efficiency was confirmed by using both qRT-PCR and immunoblotting. The sequences of shRNAs used are as follows:

NC shRNA: 5′- ACTACCGTTGTTATAGGTG -3′;

*Lars2* shRNA: 5′- CCAGAGTGGTACGGAATCAAA -3′.

### Tumor transplantation models

For primary tumor cells orthotopic transplantations, tumor cells were re-suspended in a final volume of 50μl PBS+50μl Matrigel (#354234, Corning). 1-2 × 10^6^ PyMT*^WT^*, PyMT*^TrnE^*, PyMT*^ND5^* tumor cells were injected into the fourth mammary fat pads of female WT C57BL/6J mice. The mice were euthanized 3-6 weeks after injection. For mouse cell line orthotopic transplantations, 1 × 10^6^ 4T1 cells were re-suspended and injected into the fourth mammary fat pads of female syngeneic BALB/cJ mice. The mice were euthanized 3-4 weeks after injection. Tumor size was monitored using vernier callipers with cage labels blinding from 7-14 days after tumor cell inoculation. The following formula was used to estimate tumor size (all measurements in mm): volume = (long axis) × (short axis)^2^/2. After mice were euthanized, the tumors were removed and weighed.

For analysis of the effect of the mutant microenvironment on tumors, 3 × 10^6^ PyMT*^WT^* tumor cells were injected into the fourth mammary fat pads of female mtDNA mutant *TrnE*/*ND5* mice. The mice were euthanized 3-4 weeks after injection.

For tasquinimod treatment, tasquinimod (HY-10528, MedChemExpress) was first dissolved in DMSO, then mixed with 3% sucrose (Sigma), 2% PEG300 (MedChemExpress) in PBS. Control groups received equivalent DMSO and PEG300 in PBS with 3% sucrose. Six or seventeen days after tumor cell inoculation, mice were randomly assigned and received intragastric injection of tasquinimod (10 mg/ kg) every day. In treatment experiments, mice were randomized before the start of therapy to different groups based on equal tumor size.

For MDSCs deletion, 7-8 days after tumor cell inoculation, mice were randomly assigned received intraperitoneal injection of anti-Ly6G/ Ly6C (10 mg/ kg, 1A8, Bioxcell) every other day. Cellular depletion of MDSCs was confirmed by flow cytometry.

For the metastasis assay with BALB/c mice, 1 × 10^6^ 4T1-Luc cells were injected in the fourth mammary fat pads on one flank of the mice. For *ex vivo* BLI, lung tissues carrying tumors expressing firefly luciferase were collected from sacrificed xenografted mice and incubated in 1×PBS containing 15 mg/mL D-luciferin for at least 15 min at RT to allow complete penetration of D-luciferin (ST196, Beyotime) into tissues. After washing 3 times with 1×PBS, tissues were placed on a black board and images were acquired with a PE Spectrum small animal imaging system. BLI imaging data were quantified using the living imaging software provided with the system. The color scale was shown accompanying each group of representative images.

### Human cancer data analysis

#### Database analyses

For Fig. 1a, b mtDNA single nucleotide variant (SNV) data from breast cancer patients were obtained from the Cancer Mitochondrial Atlas (TCMA) database (https://ibl.mdanderson.org/tcma/index.html) ^3^. Raw mutation data were converted into Mutation Annotation Format (MAF) files and analyzed using the online tool OmicStudio (https://www.omicstudio.cn/tool) to generate oncoplot ^47^. These plots were used to visualize the mtDNA mutation landscape across tumor samples and to identify mutation hotspots within the mitochondrial genome. Only high-confidence SNVs with annotated functional impact were included in the final analysis.

#### Pan-Cancer analyses of *Lars2*

To evaluate the expression pattern of *Lars2* across different cancer types and tissue origins, we utilized the TNMplot database (https://tnmplot.com/analysis/), which integrates gene expression data from The Cancer Genome Atlas (TCGA), the Genotype-Tissue Expression Project (GTEx), and GEO dataset. Pan-cancer expression analysis was performed to compare *Lars2* mRNA levels across diverse tumor types. In breast cancer, *Lars2* expression was further assessed across matched normal tissues, primary tumors, and metastatic lesions, demonstrating progressive upregulation during tumor advancement. Gene expression data were log2-transformed and visualized as boxplots. All analyses were conducted using the default parameters of the TNMplot platform.

#### Survival analyses

Kaplan–Meier survival curves were generated using the online tool KM Plotter (https://kmplot.com/analysis/), which integrates gene expression and clinical survival data from multiple publicly available breast cancer cohorts. Patients were stratified into high- and low-expression groups based on the median expression level of *Lars2*. Survival analysis was performed specifically within each PAM50 intrinsic molecular subtype (Normal, luminal A/B, HER2-enriched, and basal-like). The Kaplan–Meier estimator was used to assess overall survival (OS), and the log-rank test was applied to determine statistical significance between survival curves ^48^.

### Flow cytometry and analysis

#### Analysis of immune microenvironment

Single-cell suspensions were prepared from mouse tumor tissues (see **Mouse tumor dissection section**), spleens, and peripheral blood. For spleens, tissues were mechanically dissociated by grinding through a 70 μm nylon cell strainer (SORFA) in PBS, followed by red blood cell (RBC) lysis using ACK lysing buffer (#00-4300-54, Invitrogen) for 2-3 minutes at room temperature. For peripheral blood, whole blood was collected via retro-orbital into EDTA-coated tubes to prevent coagulation, and RBCs were lysed using ACK buffer. Single-cell suspensions were washed with PBS and then stained with LIVE/DEAD dye (#65-0863-14, Invitrogen) for 20 min at room temperature. Single-cell suspensions were incubated with a rat anti-mouse CD16/CD32 monoclonal antibody (#567021, BD Biosciences), followed by the surface staining on ice for 25 min. The following antibodies were used. BioLegend: CD45 (30-F11, 1:100), CD3 (17A2, 1:100), CD4 (GK1.5,1:100), CD8a (53-6.7, 1:100), CD11b (M1/70,1:100), Gr1 (RB6-8C5, 1:100), F4/80 (BM8, 1:100), Ly6G (1A8, 1:100). Immune subpopulations were determined as TAMs (CD45^+^CD11b^+^F4/80^+^), MDSCs (CD45^+^CD11b^+^Gr1^+^), CD4^+^T cells (CD45^+^CD3^+^CD4^+^), CD8^+^T cells (CD45^+^CD3^+^CD8^+^). The samples were analysed on a Flow Spectrum Analyzer (Cytek Aurora). Data analysis was done using FlowJo version 10 software.

#### Analysis of LARS2^+^ tumor cells

For intracellular detection of LARS2 in tumor-derived single-cell suspensions, surface staining was first performed. Cells were washed with PBS and incubated with anti-mouse EpCAM (1:100, G8.8, BioLegend) in staining buffer (PBS supplemented with 2% FBS and 1 mM EDTA) for 25 minutes on ice. After washing, intracellular staining of LARS2 was performed with intracellular staining kit (#00-5523-00, Invitrogen) according to the manufacturer’s protocol. In brief, cells were resuspended in Fix/Perm working solution and incubated overnight at 4 °C in the dark. The next day, permeabilized cells were washed twice with Perm buffer and incubated with primary antibody against LARS2 (1:100, 17097-1-AP, Proteintech) diluted in Perm buffer for 1h at room temperature. Afterward, cells were washed and incubated with Alexa Fluor 488-conjugated secondary antibody (1:400, #111-545-003, Jackson) for 1h at room temperature in the dark. Cells were washed twice with PBS and fluorescence signals were detected using a Cytek Aurora flow cytometer, and data were analyzed using FlowJo v10 software. EpCAM-positive cells were gated to analyze LARS2 expression within epithelial tumor cells.

#### Tumor cell sorting

For bulk mtDNA mutation analysis, LARS2^+^ tumor cells were sorted into LARS2^high^ and LARS2^low^ populations. All sorted populations were collected directly into 1.5 mL EP tubes containing lysis buffer for subsequent DNA extraction. The mtDNA mutation load was determined as described above.

#### Immune cell sorting

For immune cell sorting, single-cell suspensions were stained with LIVE/DEAD dye (#65-0863-14, Invitrogen) to exclude dead cells, followed by Fc receptor blocking with rat anti-mouse CD16/CD32 (#567021, BD Biosciences). Cells were then labeled with the following antibodies (BioLegend): CD45 (30-F11), CD3 (17A2), CD11b (M1/70), F4/80 (BM8), Ly6G (1A8), NK1.1 (PK136), CD19 (6D5). Immune subsets were sorted on a BD Fusion Cell Sorter (BD Biosciences) based on established gating strategies. For mtDNA mutation analysis, all sorted populations were collected directly into 1.5 mL EP tubes containing lysis buffer for subsequent DNA extraction. The mtDNA mutation load was determined as described above.

### Transmission Electron Microscopy (TEM)

Fresh tumor tissues (1-2 mm) were fixed in a solution of 2% paraformaldehyde and 2.5% glutaraldehyde in 0.1 M cacodylic acid buffer (pH 7.2) overnight at 4°C, then washed with phosphate buffer/cacodylate buffer. Samples were post-fixed with 1% osmium tetroxide, rinsed, and stained with 1% uranyl acetate. Dehydration was done through an ethanol series, cleared with acetone, and infiltrated with acetone/Epon 812 resin mixtures. Ultrathin sections (70 nm) were cut, stained with uranyl acetate and lead citrate, and imaged using a Talos L120C G2 transmission electron microscope at 120 kV. The size and circularity of each mitochondrion were analyzed by ImageJ.

### mtDNA Copy Number

Total DNA was extracted from tumor tissue samples using the FastPure Tissue DNA Isolation Mini Kit (DC112, Vazyme Biotech), with RNase treatment. DNA concentration was measured using a NanoDrop 2000 (Thermo), and 10 ng of DNA was used as the template for real-time quantitative PCR. The mitochondrial *Nd1* gene was normalized against the nuclear *18S* gene. Amplifications were carried out using the following primers:

mouse *Nd1* -F: 5′- gcatcttatccacgcttccg -3′,

mouse *Nd1* -R: 5′- atgtatggtggtactcccgc -3′.

mouse *18S*-F: 5′- gcaattattccccatgaacg -3′,

mouse *18S*-R: 5′- ggcctcactaaaccatccaa -3′.

### Mitochondrial functional assay

Oxygen consumption rate (OCR) and extracellular acidification rate (ECAR) were measured on a Seahorse XFe96 extracellular flux analyzer (Agilent) following the manufacturer’s instructions. In brief, cells were seeded at 2×10^4^ (PyMT*^WT^*/PyMT*^TrnE/ND5^*primary cells) cells per well in cell culture microplates (Seahorse Bioscience). After reaching 70–90% confluency, For OCR, cells were equilibrated for 1h in XF assay medium supplemented with 10 mM glucose, 2 mM sodium pyruvate and 4 mM glutamine in a non-CO_2_ incubator. OCR was monitored at baseline and throughout sequential injections of oligomycin (2 μM), FCCP (1 μM) and rotenone or antimycin A (1 μM each). For ECAR, cells were equilibrated for 1 h in XF assay medium supplemented with 4 mM glutamine in a non-CO_2_ incubator. ECAR was monitored at baseline and throughout sequential injections of glucose (10 mM), oligomycin (2 μM) and 2-DG (100 mM). Data for each well were normalized to protein concentration as determined using the Pierce BCA Protein Assay kit (Sangon Biotech) after measurement on the XFe96 machine.

### Isolation of neutrophils and Transwell assays

Single-cell suspensions were prepared from mouse bone marrow and followed by red blood cell removal using RBC lysis buffer (#00-4300-54, Invitrogen). Neutrophils were then isolated by using MojoSort™ Mouse Ly-6G Selection Kit (480123, BioLegend), following the manufacturer’s instructions. Tumor cells recruitment assay was measured using a 5-μm pore Transwell system (#3421, Corning). Primary tumor cells (PyMT*^WT^*/ PyMT*^TrnE/ND5^* Mid/High tumor cells) were isolated and was placed in the bottom of the Transwell, and 2 × 10^5^ neutrophils were added into the top chamber and placed at 37 °C, 5% CO_2_ for 12 h, then 50 μl medium was taken from the bottom wells and counted by flow cytometer. Neutrophil migration rate was calculated by dividing the number of cells entering the lower chamber by the number of cells seeded.

### Tumoroid assays

For 3D tumor spheroid culture, dissociated tumor cells were seeded at a density of 5,000 cells per well in U-bottom 96-well plates (Costar). Each well was precoated with 40 μL of growth factor-reduced Matrigel (#356234, Corning), which was solidified at 37 °C for 10 min prior to cell seeding. Cells were cultured in DMEM/F-12 medium (#11320-033, Gibco) + 10%FBS (#10091148, Gibco) + P/S antibiotics (Gibco) supplemented with EGF (10ng/uL, BD Bioscience), Y-27632 dihydrochloride (10uM, Tocris), R-spondin 1 (250 ng/mL, R&D Systems), B27 (1x, #12587010, Invitrogen) and Normocin (ant-nr-1, InvivoGen). Cultures were maintained in a humidified incubator at 37 °C and 5% CO₂.

To assess the response of tumor spheroids to doxycycline (Sigma-Aldrich, 10 μg/ ml), doxycycline was added to the culture medium 24 h after seeding. Control wells received an equivalent volume of DMSO. Spheroid growth was monitored over 4 days. Spheroid size and morphology were imaged using a Incucyte Real-Time Live Cell Analysis System, and data were analyzed using analysis software with Incucyte and ImageJ.

### H&E staining

Tumors and lungs were excised from euthanized mice and submerged in 4% PFA overnight at 4 °C, and then were transferred to 70% ethanol. Tissue embedding, slide sectioning and H&E staining were performed by the Advanced Biomedical Technology Core Facility at Westlake University. The images of the processed sections were scanned using a Panoramic (3DHISTECH) Digital Slide Scanner.

### Immunofluorescence staining

Immunofluorescence staining was performed on paraffin-embedded tumor tissue sections (4 μm thickness). After deparaffinization and rehydration through graded alcohol, antigen retrieval was performed using a sodium citrate-EDTA antigen retrieval solution (40X dilution, Beyotime) in a pressure cooker for 20 min. Sections were then blocked with 5% goat serum for 1h at room temperature to prevent nonspecific binding. For single immunofluorescence staining, sections were incubated with mouse anti-Ki67 (1:500, GB121141, Servicebio) and rabbit anti-S100A9 (1:1000, A26782PM, Abclonal) at 4 °C overnight. After washing, Alexa Fluor 488-conjugated Goat anti-Rabbit IgG (1:400, #111-545-003, Jackson) was applied for 1 h at room temperature. Nuclei were counterstained with DAPI.

For multiplex immunofluorescence staining, two different panels were performed using a TSA-based multiplex immunofluorescence kit (Servicebio) to detect multiple targets. The first panel included F4/80 (1:1000, Servicebio), Ly6G (1:1000, Servicebio), and EpCAM (1:1000, Servicebio). The second panel included RPL38 (1:500, 15055-1-AP Proteintech) and EpCAM (1:1000, Servicebio). Sections were incubated with primary antibodies at 4 °C overnight, followed by sequential incubation with species-matched HRP-conjugated secondary antibodies and TSA signal amplification using Alexa Fluor-conjugated tyramides (iF488, 555 or 647, 1:500). After each staining round, unbound antibody complexes were removed by microwave treatment to allow subsequent rounds of staining. Nuclei were counterstained with DAPI. All stained sections were imaged using the FV3000 confocal microscope (Olympus) for high-resolution whole-slide scanning. Fluorescence signal intensity and cell counts were quantified using ImageJ software.

### Human cancer tissue microarray and immunohistochemistry

Tissue microarray of human breast cancer and paired adjacent normal tissues (HLugA180Su04) were purchased from Shanghai Outdo Biotech Company (Shanghai, China). The study was approved by the Ethics Committee of Shanghai Outdo Biotech Company. Briefly, Tissue microarrays were subjected to immunohistochemical staining for LARS2. After deparaffinization, antigen retrieval, and blocking of endogenous peroxidase activity, sections were incubated with a primary antibody against LARS2 (1:1000, 17097-1-AP, Proteintech), followed by detection using a horseradish peroxidase (HRP)-conjugated secondary antibody and DAB chromogen. the sections were mounted and imaged using the Leica Aperio Scanner.

IHC-stained slides were analyzed using QuPath v0.5.1 (University of Edinburgh, UK). Tissue cores were annotated to distinguish tumor from adjacent normal regions. Mean optical intensity of LARS2 staining (mean intensity) and histo-score (H-score) were calculated for each core. H-score was determined using the formula: H-score = (1 × percentage of weakly positive cells) + (2 × percentage of moderately positive cells) + (3 × percentage of strongly positive cells), with scores ranging from 0 to 300. LARS2 expression levels in tumor and adjacent normal tissues were compared by plotting the mean intensity and H-score for each group. Representative images were selected to illustrate the staining patterns.

### Immunoblotting

Tissues and cells were pelleted and lysed using RIPA buffer (P0013B, Beyotime). Protein concentration was determined using BCA protein assay and 20-30 μg of total protein was loaded in each lane. Proteins were separated by gradient SDS-PAGE and transferred to PVDF or nitrocellulose membranes. Membranes were blocked with TBST buffer containing 5% non-fat milk for 1 h and incubated with the primary antibody in primary antibody diluent (10X, Beyotime) overnight at 4 °C: MFN2 (ABclonal A19678), DRP1 (Abcam ab184248), HSP60 (Abcam ab190828), VDAC1 (Merck MABN504), TFAM (Abcam ab131607), OXPHOS (Abcam ab110413), GAPDH (Proteintech 81640-5-RR), PLOG (Abcam ab128899), MRPS30 (Proteintech 18441-1-AP), MRPL28 (Proteintech 68467-1-Ig), RPS27 (Proteintech 15355-1-AP), LARS2 (Affinity DF13122), TOMM40 (ThermoFisher PA5-110507), TOMM20 (Cell Signaling Technology D8T4N) and β-actin (ABclonal AC026). After three washes with TBST, membranes were incubated with proper horseradish peroxidase (HRP)- or fluorescent dye-conjugated secondary antibodies (1:1000, Bioker) in TBST containing 5% non-fat milk for 1 h at room temperature. After three washes with TBST, membranes using fluorescent dye-conjugated secondary antibodies were imaged using the GEL Imaging System (AI680RGB). Relative levels were quantified by measuring band intensities on immunoblots using ImageJ software.

### qPCR and ELISA

Total RNA was extracted from tumor tissues using the FastPure Cell/Tissue Total RNA Isolation Kit (Vazyme) and reverse transcribed into cDNA using PrimeScript RT Master Mix (Takara) according to the protocol provided. RT–qPCR was performed using PowerUp SYBR Green Master Mix (Thermo Fisher Scientific) in 384-well plates. Plates were read with a Jena Bioscience Qtower384G system. Relative expression levels were normalized to the housekeeping gene *Actin*, and analyzed using qPCRsoft384 1.1 (Jena Bioscience). Amplifications were carried out using the following primers:

mouse *Actin* -F: 5′- gtgctatgttgctctagacttcg -3′,

mouse *Actin* -R: 5′- atgccacaggattccatacc -3′.

mouse *S100a9*-F: 5′- atactctaggaaggaaggacacc-3′,

mouse *S100a9*-R: 5′- tccatgatgtcatttatgagggc-3′.

Serum levels of S100A9 were measured with a commercially available ELISA kit (JHN80461, JINHENGNUO) according to the manufacturer’s instructions.

### Proteomics

#### Pretreatment

The protein sample preparation for proteomics was performed according to a previously described method ^49^. Briefly, total protein was extracted from tissues under each experimental condition using RIPA-strong buffer with 1x protease inhibitor. The samples were centrifuged at 16,000 rpm for 10 minutes at a low temperature, and the supernatant was collected. Protein concentration was determined using a BCA assay, and 25 μg of protein was used for subsequent steps. Next, 5 μL of 50 μg/μL prepared beads were added to the protein solution, followed by an equal volume of 100% ethanol. The binding mixture was incubated in a ThermoMixer at 24 °C for 10 minutes at 1,000 rpm. The unbound supernatant was then removed and discarded. The beads were washed four times with 180 μL of 80% ethanol. After washing, 100 μL of 100 mM ammonium bicarbonate containing 0.5 μg of trypsin was added, and the tube was sonicated for 30 seconds in a water bath. The digestion was incubated for 18 hours at 37 °C in a ThermoMixer at 1,000 rpm with continuous mixing. The tube was then centrifuged at 20,000 g at 24 °C for 1 minute, and the supernatant was collected. The sample was freeze-dried and resuspended in 20 μL of 0.1% formic acid (FA) for LC-MS/MS analysis.

#### LC-MS/MS Analysis

The peptides were separated by a 60 min gradient elution at a flow rate 0.300 µL/min with the Vanquish UHPLC system which is directly interfaced with the Thermo Scientific Orbitrap Exploris 480 mass spectrometer. The analytical column was a home-made fused silica capillary column (75 µm ID, 150 mm length; Upchurch, Oak Harbor, WA) packed with C-18 resin (300 A, 3 µm, Varian, Lexington, MA). Mobile phase A consisted of 0.1% formic acid, and mobile phase B consisted of 80% acetonitrile and 0.1% formic acid. The mass spectrometer was operated in the data-dependent acquisition mode using the Xcalibur 4.1 software and there is a single full-scan mass spectrum in the Orbitrap (400-1800 m/z, 60,000 resolution) followed by 20 data-dependent MS/MS scans at 30% normalized collision energy. Each mass spectrum was analyzed using the Thermo Xcalibur Qual Browser and Proteome Discovery for the database searching and quantitative analysis.

### Data processing

All analyses were performed in R (v4.3.0), where quantitative proteomic data were first filtered, normalized, and missing values imputed using the DEP package (v1.24.0) ^50^. Then the processing date were subjected to principal component analysis (PCA) using the FactoMineR package (v2.11) ^51^ to assess global differences in protein expression profiles across the five experimental groups. The differentially expressed proteins were further analyzed using the DEP package with threshold of |FC| >= 1.5 and adjusted p-value < 0.05. Functional annotation and pathway enrichment analysis were performed using ClusterProfiler package (v4.10.1) ^52^. Gene Ontology (GO) biological process (BP) terms were analyzed to identify significantly enriched pathways among differentially expressed proteins. Only terms with a false discovery rate (FDR) adjusted p-value(q-value) < 0.05, as determined by Storey, were considered statistically significant. To identify mitochondrial-enriched proteins, we cross-referenced the protein list with MitoCarta3.0, which provides a comprehensive catalog of both nuclear and mitochondrial genome-encoded mitochondrial proteins.

### Bulk RNA-seq analysis

Total RNA was extracted from 4T1 tumor xenograft tissues using TRIzol® reagent according to the manufacturer’s instructions. RNA integrity and concentration were assessed using a 5300 Bioanalyzer (Agilent) and ND-2000 NanoDrop (Thermo Fisher), respectively. High-quality RNA ([OD260/280=1.8∼2.2,OD260/230≥2.0,RQN≥6.5, 28S:18S≥1.0, >1μg] input for standard library prep) was used for library construction. Strand-specific RNA-seq libraries were prepared using Illumina® Stranded mRNA Prep Kit (San Diego, CA), involving polyA selection, mRNA fragmentation, cDNA synthesis, adapter ligation, and PCR amplification. The final libraries were sequenced on an Illumina NovaSeq X Plus platform (PE150).

Raw sequencing reads were processed using FASTQ for quality trimming and adapter removal. Clean reads were aligned to the mouse reference genome (GRCm38) using BWA-MEM (v0.7.17) (arXiv:1303.3997). Gene expression levels were quantified using FeatureCounts (v2.0.1) ^53^. Differential gene expression analysis was carried out using limma package (v3.58.1) ^54^ in R (v4.3.0) with thresholds of |log2FoldChange| > 1 and adjusted p-value < 0.05. Volcano plots were generated using the ggplot2 package (v3.5.1) to visualize significantly differentially expressed genes. Functional enrichment analysis of differentially expressed genes was conducted using ClusterProfiler package (v4.10.1) for Gene Ontology (GO) enrichment and KEGG pathway enrichment (q-value < 0.05).

### scRNA-seq

#### Sample Preparation

Single-cell suspensions were prepared from viable cells sorted from PyMT-WT/*TrnE*/*ND5* transgenic mouse models. Cells were washed, counted, and concentrated according to the 10x Genomics Cell Preparation Guide. Briefly, cell viability and concentration were assessed using a automated cell counter to ensure optimal conditions for encapsulation. Cells were suspended in appropriate buffer and loaded onto Chromium microfluidic chips for downstream barcoding and transcriptome capture.

### Library Preparation and Sequencing

Cell suspensions were loaded onto 10x Genomics Chromium microfluidic chips using 3’ v3.1 chemistry. Single-cell barcoding and partitioning were performed using the Chromium Controller (10X Genomics). Reverse transcription and library construction were carried out using the Chromium Single Cell 3’ Reagent Kit (10X Genomics) according to the manufacturer’s instructions. Libraries were sequenced on an Illumina platform (Novaseq X plus) to generate paired-end reads.

### Data Analysis

FASTQ files were generated from raw BCL data using Illumina’s standard pipeline. Quality control and preprocessing were performed using fastp and Trimmomatic, including adapter trimming, removal of low-quality bases (leading/trailing quality < 3 or average sliding window quality < 10), and filtering of reads < 26 bases. Clean reads were processed using Cell Ranger (v7.1.0) for demultiplexing, alignment to the reference genome, barcode counting, and UMI quantification. Downstream analyses were performed in R (4.3.0) using the Seurat (v4.3.0). Cells with < 200 detected genes and genes expressed in < 3 cells were filtered. Graph-based clustering (resolution = 0.6) and UMAP visualization were performed following canonical correlation analysis (CCA) integration across all samples.

Multi-group differential expression analysis among tumor subpopulations was performed using FindAllMarkers, followed by GO enrichment analysis with clusterProfiler (v3.14.0). Transcriptional regulatory networks were inferred using pySCENIC (v0.12.1) to identify cell states with distinct regulon activity. Immune composition was quantified, with emphasis on myeloid and lymphoid populations. Immunomodulatory activity was assessed using AddModuleScore in Seurat, based on gene signatures of neutrophil migration, myeloid anti-inflammatory responses, T cell activation and migration. Cell-cell communication between tumor cells and myeloid populations (macrophages and neutrophils) was analyzed using CellChat (v1.6.1) to infer receptor-ligand interactions.

### mtscATAC-seq

#### Sample preparation and library construction

Single-cell suspensions from tumors of PyMT-WT, PyMT-*TrnE* and PyMT-*ND5* mice were prepared. Live cells were sorted from different tumor-bearing mutant mice. The cells were then fixed, lysed, and permeabilized according to the Nature Protocols method ^19^. Specifically, cells were fixed in 1% paraformaldehyde at room temperature for 10 minutes, with occasional vortexing and inversion. Fixation was halted using 0.125 M glycine. The cells were then lysed on ice for 3-5 minutes using 100 μL of cold lysis buffer (10 mM Tris-HCl (pH 7.4), 10 mM NaCl, 3 mM MgCl2, 0.1% Tween-20, 0.1% Nonidet P40 Substitute, 0.01% digitonin and 1% BSA), with cell concentrations ranging from 100,000 to 300,000 cells per 100 μL. After centrifugation, cells were resuspended in a 1× dilution buffer. A 5 μL cell suspension was mixed with trypan blue for counting, and 20,000 to 30,000 cells were used for mtscATAC-seq library preparation. Libraries were prepared using the Chromium Next GEM Single Cell ATAC Kit v2, where cells were incubated in Transposition Mix at 37°C for 30 minutes. A premix of Barcoding Reagent B, Reducing Agent B, and Barcoding Enzyme was added, and the mixture was processed on a Chromium Chip E. Single-cell GEMs underwent linear amplification in a C1000 Touch Thermal Cycler with specified cycling conditions. Emulsions were coalesced, and products were purified and sequenced on an Illumina-x-plus instrument at Novogene Biotechnology Co., Ltd.

### Data processing

The nuclear mitochondrial DNA segments (NUMTs) were masked in the mm10 genome ^19^. Raw sequencing data were aligned to the modified mm10 reference genome using CellRanger-ATAC (v2.1.0). Duplicate reads were removed from the BAM file using Picard MarkDuplicates (v2.9.3) (https://broadinstitute.github.io/picard). Sinto (v0.10.1) (https://github.com/timoast/sinto) was then utilized to generate the mtscATAC-seq fragments file from the deduplicated BAM file. For the PyMT-WT, PyMT-*TrnE* and PyMT-*ND5* samples, the createArrowFiles function from the ArchR package (v1.0.2) ^55^ was used to generate arrow files for each sample, filtering out cells with a TSS enrichment score below 4 and fewer than 1000 mapped mtscATAC-seq fragments. The addDoubletScores and filterDoublets functions from the ArchR package were utilized to identify and remove potential doublet cells from each sample. The ArchRProject function from the ArchR package was then used to merge the five samples. Dimensionality reduction on the mtscATAC-seq data was performed using the addIterativeLSI function from the ArchR package. The addHarmony function in ArchR package was used for batch effect correction, and the addClusters function was applied to cluster the mtscATAC-seq data. The addUMAP function was used for Uniform Manifold Approximation and Projection (UMAP) analysis to visualize the single cells in the reduced dimension space. Finally, 23,777 high-quality cells were retained for subsequent analysis.

### Mutation load analysis

The Mitochondrial Genome Analysis Toolkit (mgatk) ^20^ was utilized to identify mutation sites and coverage in the mitochondrial genome for four mtscATAC-seq mutant samples. The coverage at the m.G14102 site/ m.G12918 site and the read count for the G>A mutation was extracted from the mgatk result files. To accurately determine the mutation load in individual cells, only cells with a coverage exceeding 10 at the m.G14102 site/ m.G12918 site were included, resulting in a total of 12,436 cells used for calculating the mutation load. The mutation load was calculated as the ratio of the read count for the G>A mutation at the m.G14102 site/ m.G12918 site to the coverage at two site. FreeBayes (v1.3.6) ^56^ was used to detect off-target sites on mtDNA, and results were shown in Extended Data Fig. 1a.

### Single cell mtDNA copy number calculation

Using mgatk ^20^, the mtDNA copy number at the single-cell level in mtscATAC-seq samples was calculated, computed as the (total bases of deduplicated reads mapped to the mtDNA genome) / (length of the mtDNA genome).

#### Functional Enrichment Analysis

The enrichGO function from the clusterProfiler package (v4.4.4) ^57^ was used to perform GO pathway enrichment analysis on the differentially expressed genes, and the top significant GO BP terms were visualized in Extended Data Fig. 2f.

### Statistics and reproducibility

Each experiment was repeated independently at least twice with similar results. Student’s t-tests were used to determine the statistical significance of differences between two groups. one-way ANOVA or two-way ANOVA was used for multiple comparison tests. Survival comparison was performed using log-rank test. The Kruskal-Wallis test followed by Dunn’s multiple comparisons test was used to assess differences in activity scores or gene expression across multiple groups in the scRNA-seq data. The following values were considered to be statistically significant: \**P* < 0.05, \*\**P* < 0.01, \*\*\**P* < 0.001, \*\*\*\**P* < 0.0001. NS, not significant. Calculations were carried out using the GraphPad Prism 10 software package. Grouped bubble plots (Fig. 4d, 4e, 5e and Extended Data Fig. 2f) and linear regression (Fig. 2c) analyses were performed using the online platform OmicStudio (https://www.omicstudio.cn/tool). All data are presented as mean ± s.e.m. No statistical method was used to predetermine sample size.

## Acknowledgements

We thank all members of the Min Jiang laboratory for insightful discussions and ongoing support. We are grateful to Pei Cai, Yue Zhang, and Bingqing Yao from the Shang Cai laboratory for technical assistance. The Westlake Laboratory Animal Resources Center is acknowledged for microinjection and mouse husbandry; the Westlake Biomedical Research Core Facilities (BRCF), Flow Cytometry Core, and High-Performance Computing Center at Westlake University for technical support; Jia Chen (Mass Spectrometry & Metabolomics Core Facility, Center for Biomedical Research Core Facilities) for proteomic analysis; and Bing Wang (Novogene Corporation Inc.) for scRNA-seq data analysis. This work was supported by the National Natural Science Foundation of China (82271897, 32570717 to M. J.), Zhejiang Provincial Natural Science Foundation of China (LZYQ25C070001 to M. J.), the Westlake University Research Center for industries of the Future (WU2022C013 to M. J.), Key R&D Program of Zhejiang (2024SSYS0033) and Westlake Education Foundation. The funders had no role in study design, data collection and analysis, decision to publish or preparation of the manuscript.

## Author contributions

M. J. conceived and led the project; N. L. conducted the majority of the experiments; G. C. conducted bioinformatics analysis of mtsc-ATAC sequencing; Q. L. conducted some animal experiments; L. Z. and J. C. constructed the mtDNA mutation mouse models; X. D. conducted bioinformatics analysis of proteomics and bulk-RNA seq; Q. Z. conducted the pre-treatment experiments for mtscATAC; X. L. assisted construct *in vitro* edited cells; R. T. provided *MMTV*-PyMT transgenic mice. Z. X. conducted mtsc-ATAC cell heterogeneity analysis; S. C. contributed to experimental design and interpretation of key results; M. J., N. L. wrote and revised the manuscript.

---

**Extended Data Fig. 1.**
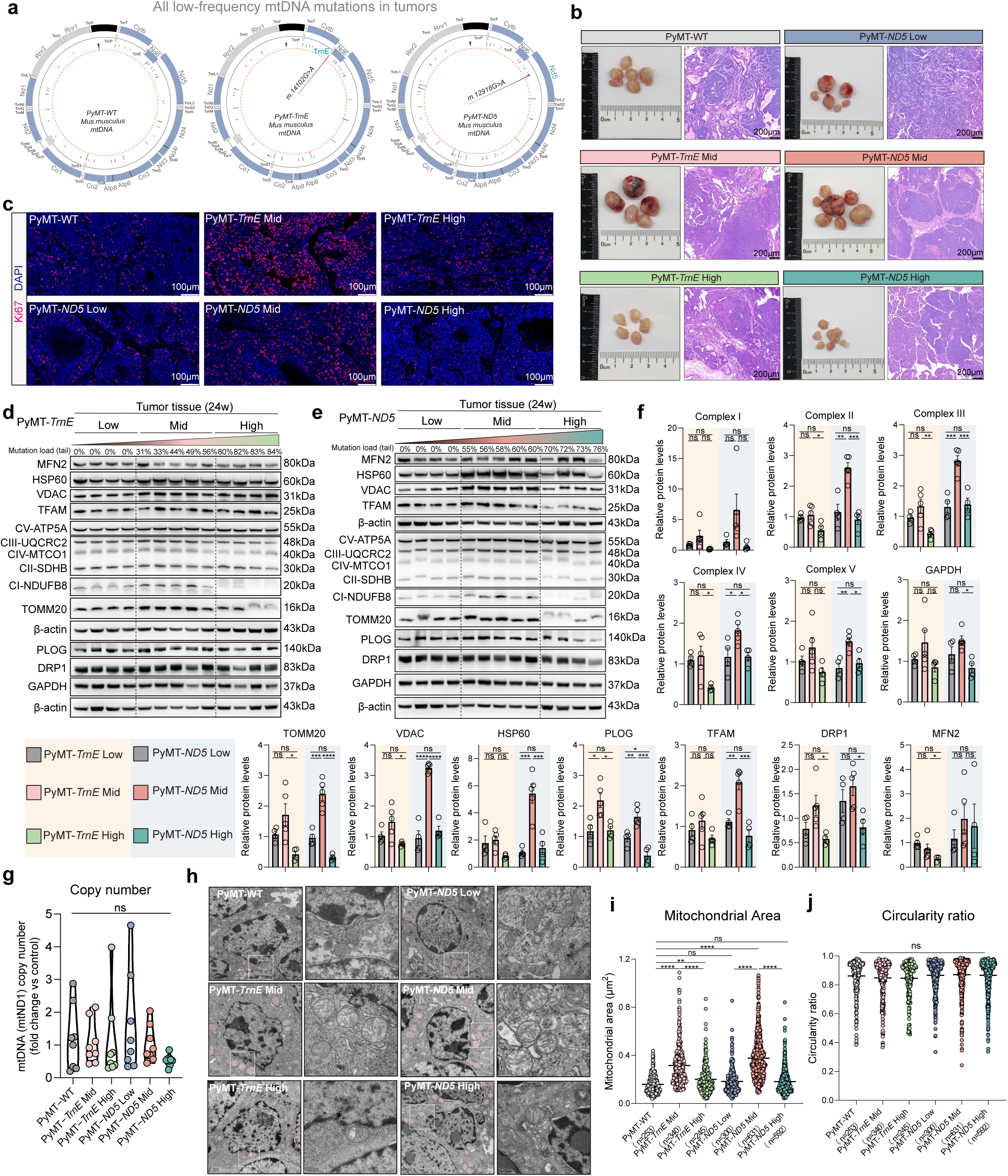
mtDNA mutation load alters mitochondrial structure and biogenesis in PyMT-*TrnE*/*ND5* breast cancer models. **a** Mitochondrial whole-genome mutation analysis of tumors from PyMT-WT, PyMT-*TrnE*, and PyMT-*ND5* mice, using bulk mtDNA data derived from mtscATAC-seq profiling. The orange dashed line indicates a 5% mutation frequency threshold. **b** Left, representative photograph of spontaneous tumors derived from PyMT-WT, PyMT-*TrnE* Mid/High, PyMT- *ND5* Low/Mid/High-mutant mice. Right, representative H&E from PyMT-WT, PyMT-*TrnE* Mid/High, PyMT-*ND5* Low/Mid/High tumors (n = 6 mice per group). Scale bars (H&E images), 200 μm. **c** Representative Ki67 staining images of tumors from PyMT-WT, PyMT-*TrnE* Mid/High, and PyMT-*ND5* Low/Mid/High groups (n = 6 mice per group). Scale bars, 100 μm. **d-f** Immunoblots showing mitochondria biogenesis related proteins expression in PyMT-*TrnE* (**d**) and PyMT-*ND5* (**e**) bulk tumors and statistics showing expression of Complex I-V, GAPDH, DRP1, MFN2, PLOG, TOMM20, VDAC, HSP60 in PyMT-*TrnE*/*ND5* Low, Mid and High groups (**f**). β-Actin is a loading control. n = 4-5 biological replicates. **g** mtDNA copy number of bulk tumors in PyMT-WT and PyMT-*TrnE*/*ND5* mice (n = 8 mice per group). **h-j** Representative mitochondrial ultrastructural images from tumor cells in PyMT-WT, PyMT-*TrnE* Mid/ High, PyMT- *ND5* Low/Mid/ High mice (**h**) (n = 3 mice per group). Quantification of relative mitochondrial area (**i**) and circularity ratio of mitochondria (**j**) in tumors across groups. n = 253, 340, 245, 300, 631 and 592 mitochondria for PyMT-WT, PyMT-*TrnE* Mid/ High, PyMT-*ND5* Low/Mid/ High group, respectively. Data are as mean ± s.e.m. One-way ANOVA with Tukey’s multiple comparison test (**f, g, i, j**). NS, not significant; \**P* < 0.05; \*\**P* < 0.01; \*\*\**P* < 0.001; \*\*\*\**P* < 0.0001.

**Extended Data Fig. 2.**
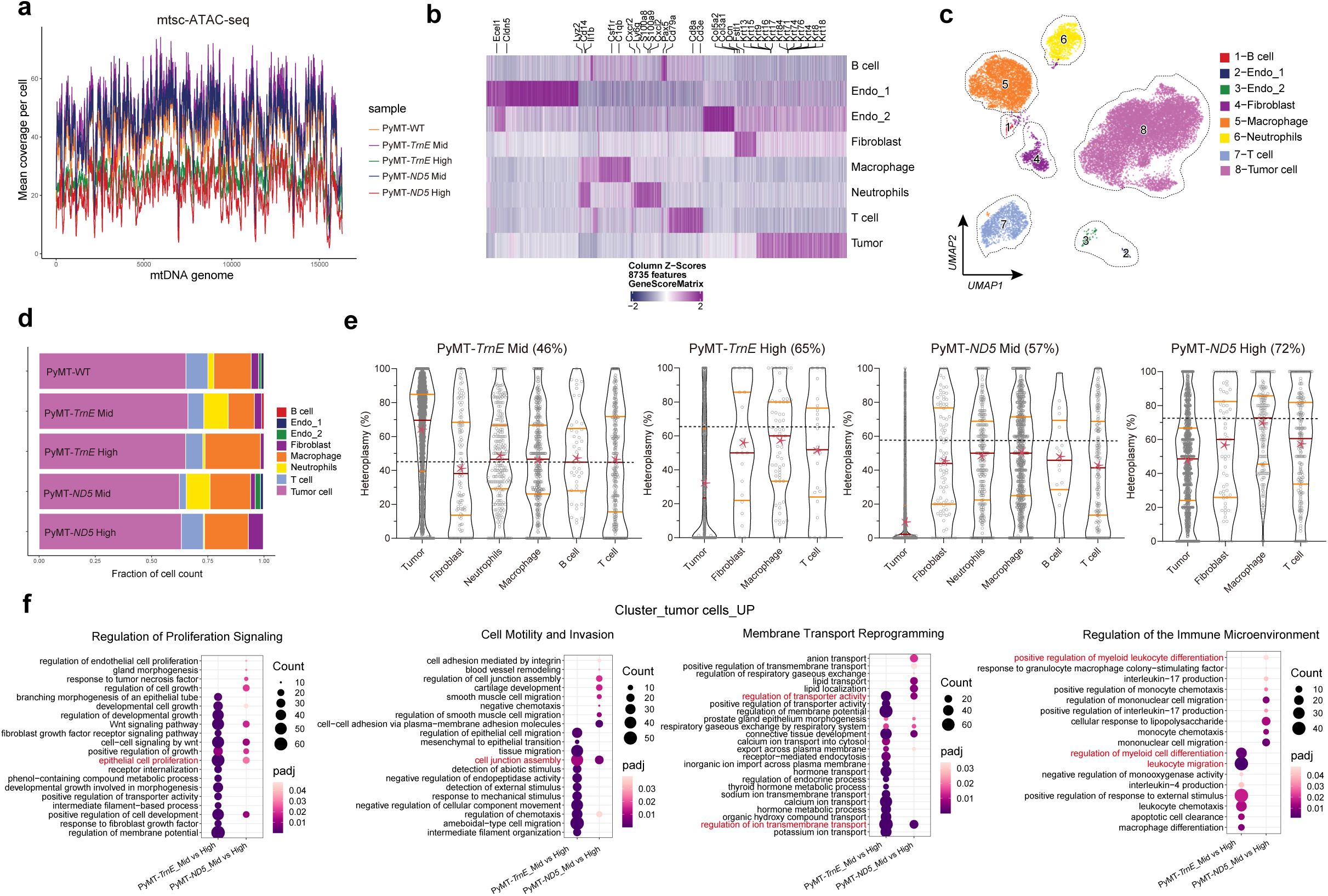
mtscATAC-seq reveals intratumoral heterogeneity of mtDNA mutations across distinct cell populations in the tumor microenvironment. **a** Visualization of mitochondrial genome coverage for five mtscATAC-seq samples. The mean coverage per cell is shown. **b** Heatmap showing expression of marker genes identified from mtscATAC-seq clusters, used to define and annotate distinct cell populations. **c** Reduced dimensionality projection and clustering of mtscATAC-seq samples from tumors of PyMT-WT, PyMT-*TrnE* Mid/ High, PyMT-*ND5* Mid/ High group. Major cell types and clusters are annotated. **d** The relative proportions of various cell types in the tumor microenvironment across PyMT-WT, PyMT-*TrnE* Mid/ High, PyMT- *ND5* Mid/ High tumor samples. **e** Heteroplasmy distribution across multiple cell types from tumors of PyMT-*TrnE* Mid/High, PyMT-*ND5* Mid/High group. Each dot represents the mutation load of a single cell (read coverage at m.G14102 site/ m.G12918 site ≥10). Median (red line), quartile range (orange line), and mean (red asterisk) are indicated. **f** Bubble plot showing differentially enriched GO pathways in PyMT-*TrnE* Mid vs High and PyMT-*ND5* Mid vs High comparisons. The enriched pathways are grouped into manually curated categories, including "Regulation of Proliferation Signaling" (top left), "Cell Motility and Invasion" (top right), "Membrane Transport Reprogramming" (bottom left), and "Regulation of the Immune Microenvironment" (bottom right). Bubble size represents the number of enriched genes (count), and color intensity indicates the adjusted p-value (padj).

**Extended Data Fig. 3.**
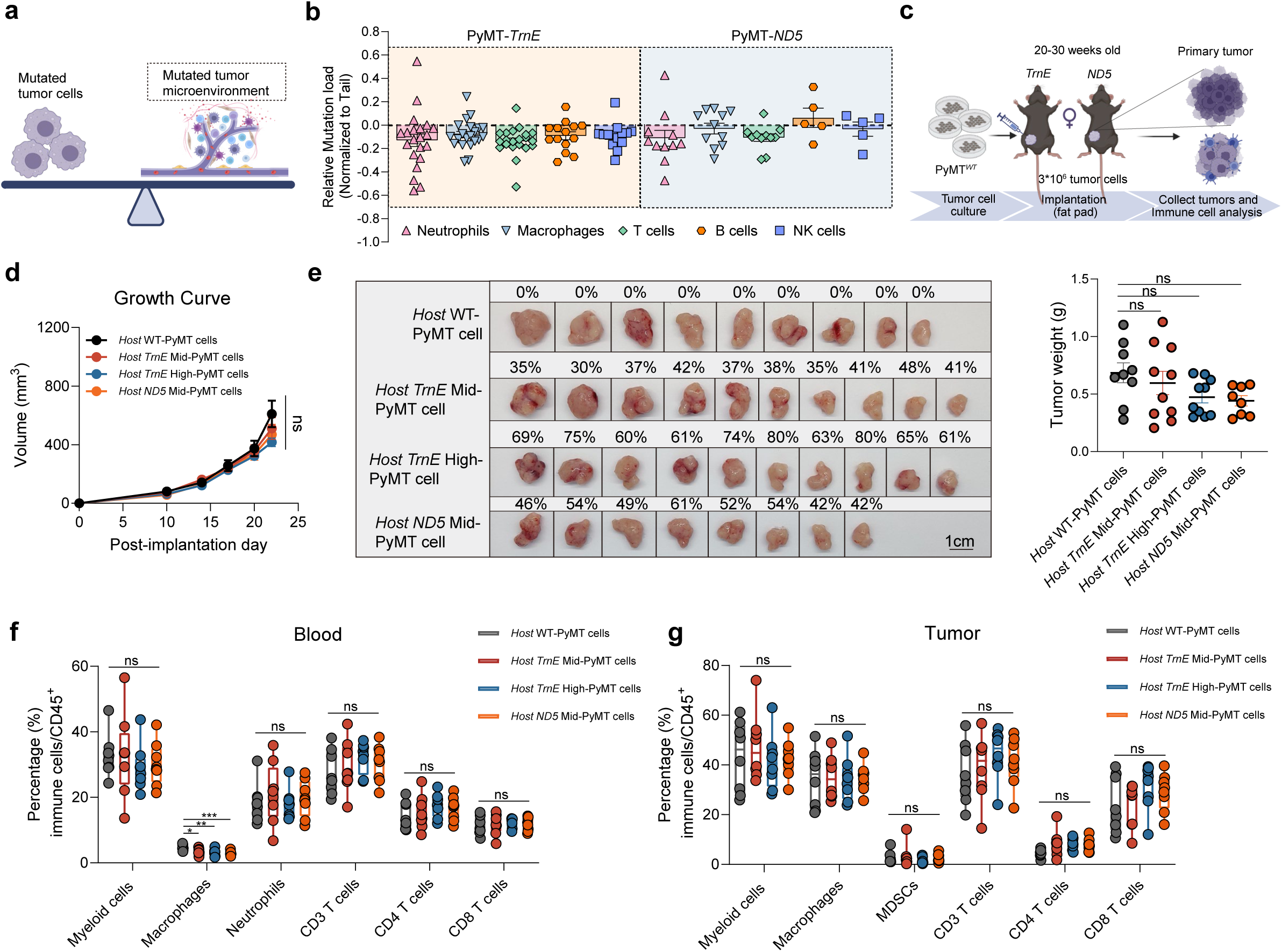
The mtDNA-mutant tumor microenvironment does not significantly alter tumor growth or immune composition. **a** Schematic showing the balance between mutant tumor cells and other mutated cells in the tumor microenvironment. **b** Mutation load analysis of sorted immune cells (neutrophils, macrophages, T cells, B cells, NK cells) from PyMT-*TrnE*/*ND5* mice tumors, normalized to the tail-derived mtDNA mutation level of the same mouse (PyMT-*TrnE*: neutrophils, n = 21; macrophages, n = 21; T cells, n = 21; B cells, n = 14; NK cells, n = 14; PyMT-*ND5*: neutrophils, n = 11; macrophages, n = 11; T cells, n = 11; B cells, n = 5; NK cells, n = 5). **c** Engraftment strategy for implanting PyMT*^WT^* primary tumor cells into *TrnE*/*ND5* mtDNA mutant mice. **d, e** PyMT*^WT^* tumor growth in *TrnE*/*ND5* mtDNA mutant mice (**d**). Tumor weight (**e**, right) and images (**e**, left) of PyMT*^WT^* tumors grown in *TrnE*/*ND5* mtDNA mutant mice. (*Host* WT-PyMT cells, n = 9; *Host TrnE* Mid-PyMT cells, n = 10; *Host TrnE* High-PyMT cells, n = 10; *Host ND5* Mid-PyMT cells, n = 8). Scale bars, 1 cm. **f, g** The percentage of indicated immune cells in the blood (**f**) and tumor (**g**) of *TrnE*/*ND5* tumor-bearing mice engrafted with PyMT*^WT^* cells as determined by flow cytometry (n = 8 mice per group). Data are as mean ± s.e.m. two-way ANOVA (**d**) and One-way ANOVA with Tukey’s multiple comparison test (**e, f, g**). NS, not significant; \**P* < 0.05; \*\**P* < 0.01; \*\*\**P* < 0.001.

**Extended Data Fig. 4.**
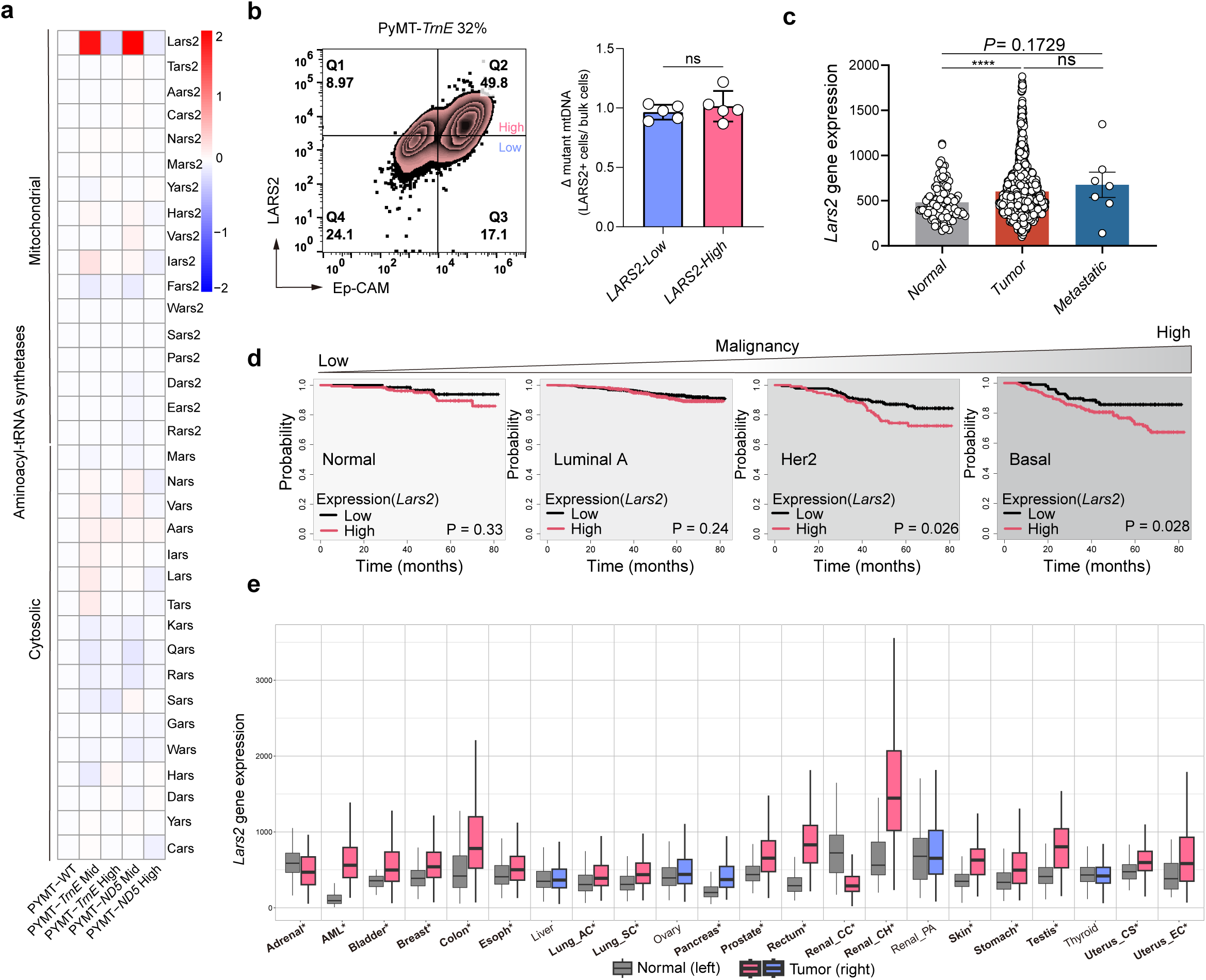
Pan-cancer analysis reveals LARS2 as a prognostic marker associated with mtDNA mutation status. **a** Heatmap showing expression of mitochondrial and cytosolic aminoacyl-tRNA synthetase (ARS) in tumor cell clusters from PyMT-WT, PyMT-*TrnE* Mid/High, and PyMT-*ND5* Mid/High groups based on scRNA-seq data. **b** Representative flow cytometry plots (left) from a PyMT-*TrnE* 32% sample show gating strategy for Ep-CAM^+^LARS2^+^ cells, further stratified into LARS2-low and LARS2-high subpopulations based on LARS2 signal intensity. Bar plot (right) compares mutation load between FACS-sorted LARS2-low and LARS2-high fractions. n = 5 biological replicates. **c** Expression of *Lars2* mRNA across normal, primary tumor, and metastatic tissues based on integrated analysis of multiple cancer databases (TNMplot.com) (Normal, n=113; Tumor, n=1097; Metastatic, n=7). **d** Kaplan-Meier analysis showing the overall survival of patients with breast cancer stratified by PAM50 subtype (Normal, Luminal A, Her2, Basal) and *Lars2* expression status (Low vs High). **e** Expression of *Lars2* mRNA in human cancers versus corresponding normal tissues from the multiple database (TNMplot.com). Adrenocortical Carcinoma (Adrenal), Acute Myeloid Leukemia (AML), Bladder Urothelial Carcinoma (Bladder), Breast Invasive Carcinoma (Breast), Colon Adenocarcinoma (Colon), Esophageal Carcinoma (Esoph), Liver Hepatocellular Carcinoma (Liver), Lung Adenocarcinoma (Lung_AC), Lung Squamous Cell Carcinoma (Lung_SC), Ovarian Serous Cystadenocarcinoma (Ovary), Pancreatic Adenocarcinoma (Pancreas), Prostate Adenocarcinoma (Prostate), Rectum Adenocarcinoma (Rectum), Kidney Clear Cell Carcinoma (Renal_CC), Kidney Chromophobe Carcinoma (Renal_CH), Kidney Papillary Adenocarcinoma (Renal_PA), Skin Cutaneous Melanoma (Skin), Stomach Adenocarcinoma (Stomach), Testicular Germ Cell Tumors (Testis), Thyroid Carcinoma (Thyroid), Uterine Carcinosarcoma (Uterus_CS), and Uterine Endometrial Carcinoma (Uterus_EC). Red: significance (*P* < 0.05); blue: no significance (*P*≥0.05). Data are as mean ± s.e.m. Student’s two-sided unpaired t-test **(b),** Kruskal-Wallis test **(c),** log-rank test **(d),** and Wilcoxon rank-sum test **(e)**. NS, not significant; \**P* < 0.05; \*\*\*\**P* < 0.0001.

**Extended Data Fig. 5.**
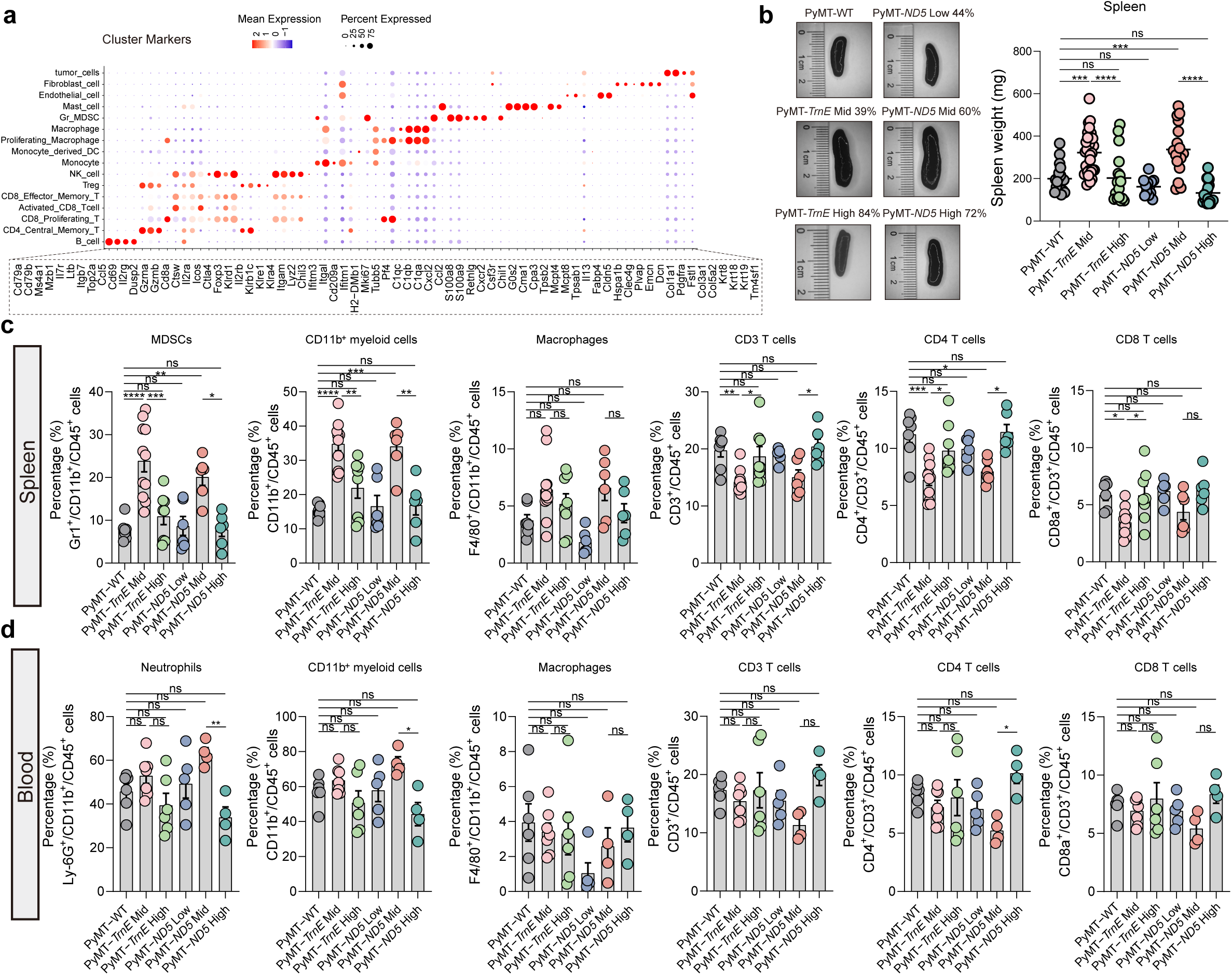
mtDNA heteroplasmy drives systemic expansion of myeloid cells in a mutation load–dependent manner. **a** Gene expression from scRNA-seq experiment characterizing expression of lineage-defining genes in cell clusters. **b** Representative spleen images (left) and quantification (right) of spleen mass from PyMT-WT, PyMT-*TrnE*/*ND5* mice. n = 18, 30, 16, 12, 17 and 18 mice for PyMT-WT, PyMT-*TrnE* Mid/ High, PyMT-*ND5* Low/Mid/ High group, respectively. **c** The percentage of indicated immune cell types in the spleen of PyMT-WT, PyMT-*TrnE*, PyMT- *ND5* mice as determined by flow cytometry. n = 7, 11, 8, 6, 6 and 6 mice for PyMT-WT, PyMT-*TrnE* Mid/ High, PyMT-*ND5* Low/Mid/ High group, respectively. **d** The percentage of indicated immune cell types in the blood of PyMT-WT, PyMT-*TrnE*, PyMT-*ND5* mice as determined by flow cytometry. n = 6, 7, 6, 5, 4 and 4 mice for PyMT-WT, PyMT-*TrnE* Mid/ High, PyMT-*ND5* Low/Mid/ High group, respectively. Data are as mean ± s.e.m. One-way ANOVA with Tukey’s multiple comparison test (**b, c, d**). NS, not significant; \**P* < 0.05; \*\**P* < 0.01; \*\*\**P* < 0.001; \*\*\*\**P* < 0.0001.

**Extended Data Fig. 6.**
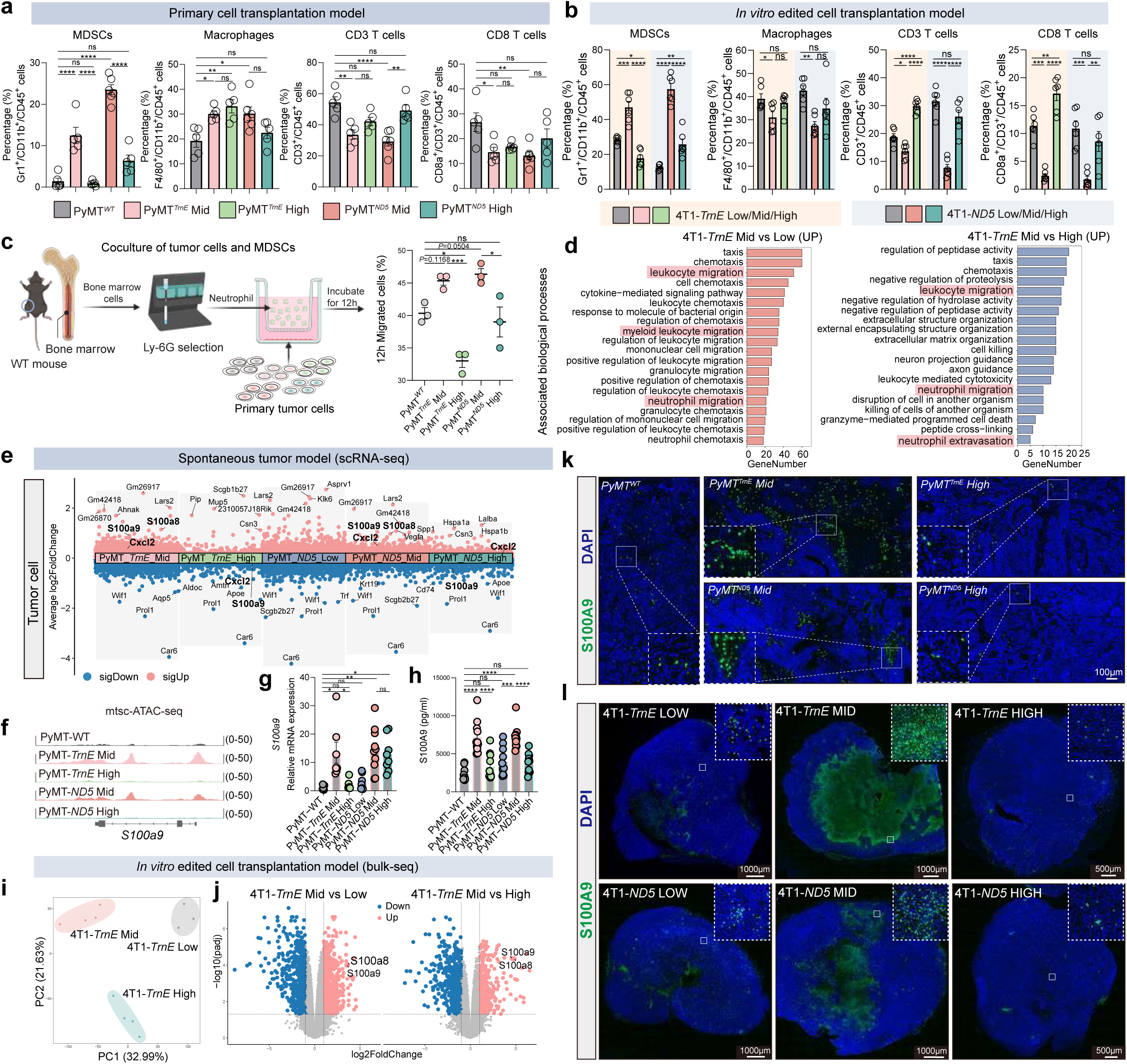
Moderate-mutant tumor cells drive S100A8/A9–dependent MDSCs recruitment and pro-metastatic immune remodeling. **a** The percentage of indicated immune cell types in orthotopic tumors derived from PyMT*^WT^*, PyMT*^TrnE^* Mid/High and PyMT*^ND5^* Mid/High cell inoculations in wild-type mice as determined by flow cytometry (n = 5-6 mice per group). **b** The percentage of indicated immune cell types in orthotopic tumors derived from 4T1-*TrnE/ND5*Low, Mid, and High cell inoculations in wild-type mice as determined by flow cytometry (n = 6 mice per group). **c** Schematic (left) showing the co-culture strategy of primary PyMT*^WT^*, PyMT*^TrnE^* Mid/High and PyMT*^ND5^* Mid/High tumor cells with purified Ly6G^+^ neutrophils isolated from bone marrow of wild-type mice. Bar plot (right) showing the percentage of Ly6G^+^ neutrophils that migrated to the lower chamber after 12 hours. n = 3 biological replicates. **d** GO pathway enrichment analysis showing upregulated biological processes in 4T1-*TrnE* Mid vs Low (red) and 4T1-*TrnE* Mid vs High (blue) comparisons, based on bulk RNA-seq data from bulk tumors derived from 4T1-*TrnE* Low, Mid, and High cell inoculations (4T1-*TrnE* Low (n = 3), Mid (n = 4), and High (n = 4)). **e** Differentially expressed genes in tumor cells clusters from PyMT-*TrnE* Mid/High and PyMT-*ND5* Low/Mid/High groups compared to PyMT-WT, based on scRNA-seq analysis (Blue: down, red: up). The y-axis represents average log2 fold change, and S100A8, S100A9, and CXCL2 are indicated in bold. **f** Genome browser visualization showing S100A9 locus accessibility across PyMT-WT, PyMT-*TrnE* Mid/High, and PyMT-*ND5* Mid/High groups, based on pre-clustering mtsc-ATAC-seq data. **g** mRNA expression levels of S100A9 in bulk tumors from PyMT-WT, PyMT-*TrnE*/*ND5* Low/Mid/High groups, as determined by qPCR, n = 6, 7, 6, 8, 10 and 8 mice for PyMT-WT, PyMT-*TrnE* Mid/ High PyMT-*ND5* Low/Mid/ High group, respectively. **h** ELISA quantification of serum S100A9 levels in PyMT-WT, PyMT-*TrnE* Mid/High, and PyMT-*ND5* Low/Mid/High breast cancer mice (n = 11 mice per group). **i, j** Transcriptomic differences among bulk tumors derived from 4T1-*TrnE* Low, Mid, and High cell inoculations by PCA **(i)**, Each circle represents one mouse. Volcano plots depict differentially expressed genes in 4T1-*TrnE* Mid vs Low (**j**, left) and 4T1-*TrnE* Mid vs High (**j**, right) (Blue: down, red: up). S100A8 and S100A9 are indicated (4T1-*TrnE* Low (n = 3), Mid (n = 4), and High (n = 4)). **k** Representative fluorescence images showing S100A9 signal in tumors from PyMT*^WT^*, PyMT*^TrnE^* Mid/High and PyMT*^ND5^* Mid/High groups (n = 3 independent tumor sections per group). Scale bars, 100 μm. **l** Representative fluorescence images showing S100A9 signal in tumors from 4T1-*TrnE*/*ND5* Low, Mid, and High groups (n = 3 independent tumor sections per group). Scale bars are shown in individual panels. Data are as mean ± s.e.m. One-way ANOVA with Tukey’s multiple comparison test (**a, b, c, g, h**). Schematic in **c** were created in BioRender. NS, not significant; \**P* < 0.05; \*\**P* < 0.01; \*\*\**P* < 0.001; \*\*\*\**P* < 0.0001.

**Extended Data Fig. 7.**
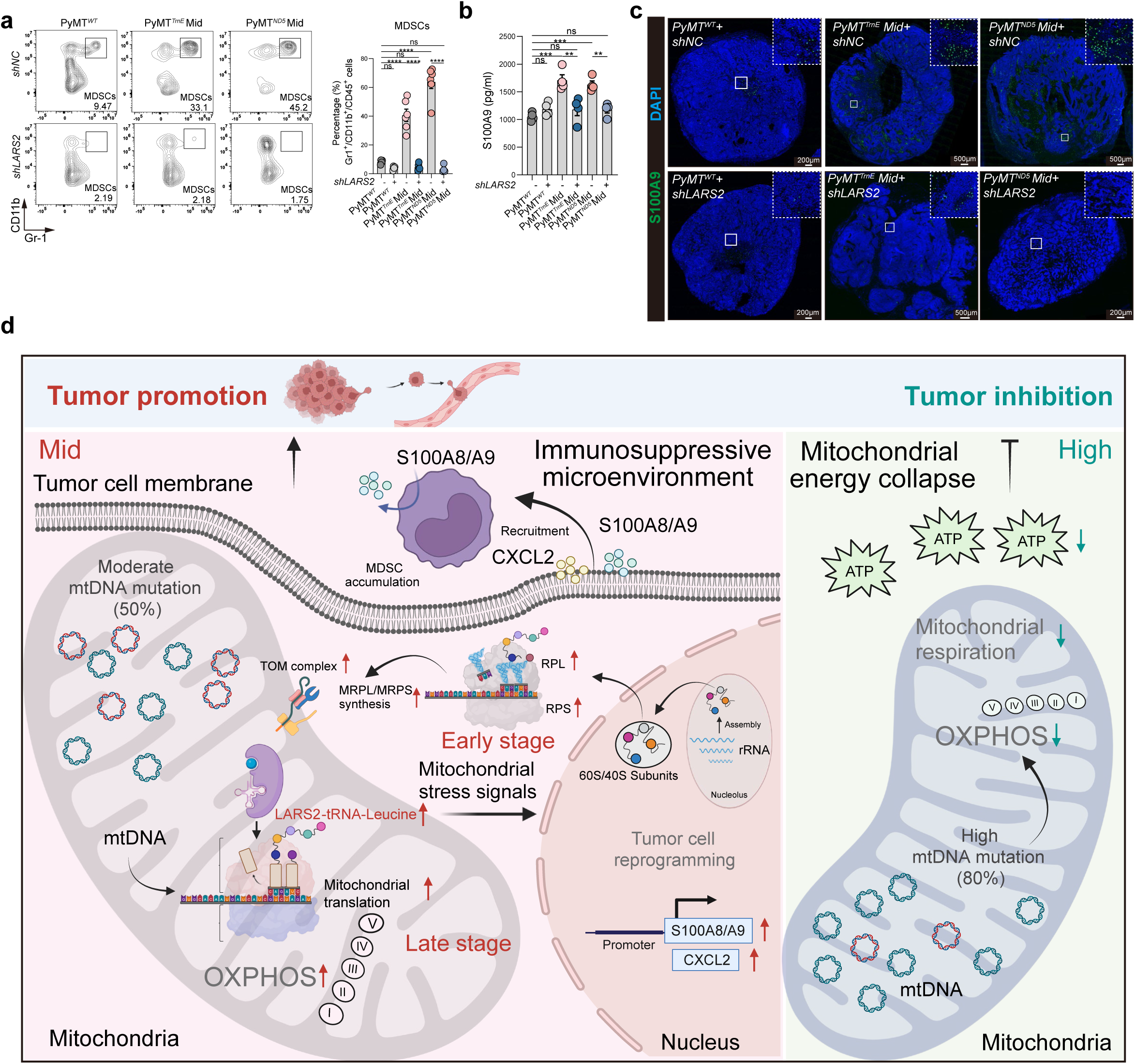
LARS2 knockdown suppresses S100A9 secretion and MDSCs infiltration in moderate-heteroplasmy tumors. **a** Representative flow cytometry plots (left) showing the proportion of MDSCs in tumors from *Lars2* non-knockdown and knockdown groups of PyMT*^WT^*, PyMT*^TrnE^* Mid, and PyMT*^ND5^* Mid. Bar plots (right) showing the percentage of MDSCs across groups (n = 4-6 tumors per group). **b** ELISA quantification of serum S100A9 levels in mice bearing *Lars2* non-knockdown and knockdown PyMT*^WT^*, PyMT*^TrnE^*Mid, and PyMT*^ND5^* Mid tumors (n = 4 mice per group). **c** Representative fluorescence images showing S100A9 signal in tumors from *Lars2* non-knockdown and knockdown PyMT*^WT^*, PyMT*^TrnE^*Mid, and PyMT*^ND5^* Mid groups (n = 3 independent tumor sections per group). Scale bars are shown in individual panels. **d** Schematic model illustrating how mtDNA heteroplasmy level dictates tumor fate. Moderate mtDNA mutations enhance mitochondrial translation through upregulation of LARS2 and mitochondrial ribosomal genes, leading to increased mitochondrial protein synthesis. This adaptive shift promotes the secretion of S100A8/A9 and CXCL2, which recruit myeloid-derived suppressor cells (MDSCs) and establish an immunosuppressive tumor microenvironment. Mitochondrial translation activation and immune evasion drive tumor growth and metastasis. In contrast, high mutations result in respiratory dysfunction and metabolic collapse, which impair tumor fitness and immune signaling, leading to attenuated progression. Schematic was created in BioRender. Data are as mean ± s.e.m. One-way ANOVA with Tukey’s multiple comparison test (**a, b**). Schematic in **d** was created in BioRender. NS, not significant; \*\**P* < 0.01; \*\*\**P* < 0.001; \*\*\*\**P* < 0.0001.

## References

1 Gustafsson, C. M., Falkenberg, M. & Larsson, N. G. Maintenance and Expression of Mammalian Mitochondrial DNA. Annu Rev Biochem 85, 133–160, doi:10.1146/annurev-biochem-060815-014402 (2016).

2 Kauppila, T. E. S., Kauppila, J. H. K. & Larsson, N. G. Mammalian Mitochondria and Aging: An Update. Cell Metab 25, 57–71, doi:10.1016/j.cmet.2016.09.017 (2017).

3 Yuan, Y. et al. Comprehensive molecular characterization of mitochondrial genomes in human cancers. Nat Genet 52, 342–352, doi:10.1038/s41588-019-0557-x (2020).

4 Kopinski, P. K., Singh, L. N., Zhang, S., Lott, M. T. & Wallace, D. C. Mitochondrial DNA variation and cancer. Nat Rev Cancer 21, 431–445, doi:10.1038/s41568-021-00358-w (2021).

5 Barrera-Paez, J. D. & Moraes, C. T. Mitochondrial genome engineering coming-of-age. Trends Genet 38, 869–880, doi:10.1016/j.tig.2022.04.011 (2022).

6 Kim, M., Mahmood, M., Reznik, E. & Gammage, P. A. Mitochondrial DNA is a major source of driver mutations in cancer. Trends in Cancer 8, 1046–1059, doi:10.1016/j.trecan.2022.08.001 (2022).

7 Gopal, R. K. et al. Early loss of mitochondrial complex I and rewiring of glutathione metabolism in renal oncocytoma. Proc Natl Acad Sci U S A 115, E6283–E6290, doi:10.1073/pnas.1711888115 (2018).

8 Mahmood, M. et al. Mitochondrial DNA mutations drive aerobic glycolysis to enhance checkpoint blockade response in melanoma. Nat Cancer 5, 659–672, doi:10.1038/s43018-023-00721-w (2024).

9 Gorelick, A. N. et al. Respiratory complex and tissue lineage drive recurrent mutations in tumour mtDNA. Nat Metab 3, 558–570, doi:10.1038/s42255-021-00378-8 (2021).

10 Stewart, J. B. & Chinnery, P. F. The dynamics of mitochondrial DNA heteroplasmy: implications for human health and disease. Nat Rev Genet 16, 530–542, doi:10.1038/nrg3966 (2015).

11 Parakatselaki, M. E. & Ladoukakis, E. D. mtDNA Heteroplasmy: Origin, Detection, Significance, and Evolutionary Consequences. Life (Basel) 11, doi:10.3390/life11070633 (2021).

12 Gaude, E. et al. NADH Shuttling Couples Cytosolic Reductive Carboxylation of Glutamine with Glycolysis in Cells with Mitochondrial Dysfunction. Mol Cell 69, 581–593 e587, doi:10.1016/j.molcel.2018.01.034 (2018).

13 Shelton, S. D. et al. Pathogenic mitochondrial DNA mutations inhibit melanoma metastasis. Sci Adv 10, eadk8801, doi:10.1126/sciadv.adk8801 (2024).

14 Smith, A. L. et al. Age-associated mitochondrial DNA mutations cause metabolic remodelling that contributes to accelerated intestinal tumorigenesis. Nat Cancer 1, 976–989, doi:10.1038/s43018-020-00112-5 (2020).

15 Li-Harms, X. et al. Somatic mtDNA mutation burden shapes metabolic plasticity in leukemogenesis. Sci Adv 11, eads8489, doi:10.1126/sciadv.ads8489 (2025).

16 Mok, B. Y. et al. A bacterial cytidine deaminase toxin enables CRISPR-free mitochondrial base editing. Nature 583, 631–637, doi:10.1038/s41586-020-2477-4 (2020).

17 Ru, Y. et al. Maternal age enhances purifying selection on pathogenic mutations in complex I genes of mammalian mtDNA. Nat Aging 4, 1211–1230, doi:10.1038/s43587-024-00672-6 (2024).

18 Zhang, L. et al. Age-dependent accumulation of mitochondrial tRNA mutations in mouse kidneys linked to mitochondrial kidney diseases. Nat Aging 5, 1317–1339, doi:10.1038/s43587-025-00909-y (2025).

19 Lareau, C. A. et al. Mitochondrial single-cell ATAC-seq for high-throughput multi-omic detection of mitochondrial genotypes and chromatin accessibility. Nat Protoc 18, 1416–1440, doi:10.1038/s41596-022-00795-3 (2023).

20 Lareau, C. A. et al. Massively parallel single-cell mitochondrial DNA genotyping and chromatin profiling. Nat Biotechnol 39, 451–461, doi:10.1038/s41587-020-0645-6 (2021).

21 Tzagoloff, A., Akai, A., Kurkulos, M. & Repetto, B. Homology of yeast mitochondrial leucyl-tRNA synthetase and isoleucyl- and methionyl-tRNA synthetases of Escherichia coli. J Biol Chem 263, 850–856 (1988).

22 van Vlerken-Ysla, L., Tyurina, Y. Y., Kagan, V. E. & Gabrilovich, D. I. Functional states of myeloid cells in cancer. Cancer Cell 41, 490–504, doi:10.1016/j.ccell.2023.02.009 (2023).

23 Chen, Y., Ouyang, Y., Li, Z., Wang, X. & Ma, J. S100A8 and S100A9 in Cancer. Biochim Biophys Acta Rev Cancer, 188891, doi:10.1016/j.bbcan.2023.188891 (2023).

24 Luo, W. et al. S100A8/A9 perturbation in bone marrow blunts antitumor immunity by promoting protumorigenic myelopoiesis in mouse models. Sci Transl Med 17, eadr3963, doi:10.1126/scitranslmed.adr3963 (2025).

25 Zhang, R. et al. PMN-MDSCs modulated by CCL20 from cancer cells promoted breast cancer cell stemness through CXCL2-CXCR2 pathway. Signal Transduct Target Ther 8, 97, doi:10.1038/s41392-023-01337-3 (2023).

26 Russell, O. M., Gorman, G. S., Lightowlers, R. N. & Turnbull, D. M. Mitochondrial Diseases: Hope for the Future. Cell 181, 168–188, doi:10.1016/j.cell.2020.02.051 (2020).

27 Larsson, N. G. & Wedell, A. Mitochondria in human disease. J Intern Med 287, 589–591, doi:10.1111/joim.13088 (2020).

28 Wallace, D. C. & Chalkia, D. Mitochondrial DNA genetics and the heteroplasmy conundrum in evolution and disease. Cold Spring Harb Perspect Biol 5, a021220, doi:10.1101/cshperspect.a021220 (2013).

29 Suomalainen, A. & Battersby, B. J. Mitochondrial diseases: the contribution of organelle stress responses to pathology. Nat Rev Mol Cell Biol 19, 77–92, doi:10.1038/nrm.2017.66 (2018).

30 Ratnaike, T. E. et al. MitoPhen database: a human phenotype ontology-based approach to identify mitochondrial DNA diseases. Nucleic Acids Res 49, 9686–9695, doi:10.1093/nar/gkab726 (2021).

31 Chen, L. et al. A mitochondrial disease model is generated and corrected using engineered base editors in rat zygotes. Nat Biotechnol, doi:10.1038/s41587-025-02684-y (2025).

32 Gammage, P. A. et al. Genome editing in mitochondria corrects a pathogenic mtDNA mutation in vivo. Nat Med 24, 1691–1695, doi:10.1038/s41591-018-0165-9 (2018).

33 Bacman, S. R. et al. MitoTALEN reduces mutant mtDNA load and restores tRNA(Ala) levels in a mouse model of heteroplasmic mtDNA mutation. Nat Med 24, 1696–1700, doi:10.1038/s41591-018-0166-8 (2018).

34 Boscenco, S. et al. Functionally dominant hotspot mutations of mitochondrial ribosomal RNA genes in cancer. Nat Genet, doi:10.1038/s41588-025-02374-0 (2025).

35 Hornig-Do, H. T. et al. Human mitochondrial leucyl tRNA synthetase can suppress non cognate pathogenic mt-tRNA mutations. EMBO Mol Med 6, 183–193, doi:10.1002/emmm.201303202 (2014).

36 Wang, Z. et al. Leucine-tRNA-synthase-2-expressing B cells contribute to colorectal cancer immunoevasion. Immunity 55, 1067–1081 e1068, doi:10.1016/j.immuni.2022.04.017 (2022).

37 Ross-Inta, C., Tsai, C. Y. & Giulivi, C. The mitochondrial pool of free amino acids reflects the composition of mitochondrial DNA-encoded proteins: indication of a post-translational quality control for protein synthesis. Biosci Rep 28, 239–249, doi:10.1042/BSR20080090 (2008).

38 Li, Q. & Hoppe, T. Role of amino acid metabolism in mitochondrial homeostasis. Front Cell Dev Biol 11, 1127618, doi:10.3389/fcell.2023.1127618 (2023).

39 Veglia, F., Sanseviero, E. & Gabrilovich, D. I. Myeloid-derived suppressor cells in the era of increasing myeloid cell diversity. Nat Rev Immunol 21, 485–498, doi:10.1038/s41577-020-00490-y (2021).

40 Li, K. et al. Myeloid-derived suppressor cells as immunosuppressive regulators and therapeutic targets in cancer. Signal Transduct Target Ther 6, 362, doi:10.1038/s41392-021-00670-9 (2021).

41 Shi, T. et al. CHI3L3(+) immature neutrophils inhibit anti-tumor immunity and impede immune checkpoint blockade therapy in bone metastases. Cancer Cell, doi:10.1016/j.ccell.2025.07.007 (2025).

42 Garner, H. et al. Understanding and reversing mammary tumor-driven reprogramming of myelopoiesis to reduce metastatic spread. Cancer Cell 43, 1279–1295 e1279, doi:10.1016/j.ccell.2025.04.007 (2025).

43 Adrover, J. M. et al. Neutrophils drive vascular occlusion, tumour necrosis and metastasis. Nature, doi:10.1038/s41586-025-09278-3 (2025).

44 Fu, A. et al. Tumor-resident intracellular microbiota promotes metastatic colonization in breast cancer. Cell, doi:10.1016/j.cell.2022.02.027 (2022).

45 Liu, Q. et al. Hi-TOM: a platform for high-throughput tracking of mutations induced by CRISPR/Cas systems. Sci China Life Sci 62, 1–7, doi:10.1007/s11427-018-9402-9 (2019).

46 Sun, T., Liu, Q., Chen, X., Hu, F. & Wang, K. Hi-TOM 2.0: an improved platform for high-throughput mutation detection. Sci China Life Sci 67, 1532–1534, doi:10.1007/s11427-024-2555-x (2024).

47 Mayakonda, A., Lin, D. C., Assenov, Y., Plass, C. & Koeffler, H. P. Maftools: efficient and comprehensive analysis of somatic variants in cancer. Genome Res 28, 1747–1756, doi:10.1101/gr.239244.118 (2018).

48 Gyorffy, B. Survival analysis across the entire transcriptome identifies biomarkers with the highest prognostic power in breast cancer. Comput Struct Biotechnol J 19, 4101–4109, doi:10.1016/j.csbj.2021.07.014 (2021).

49 Hughes, C. S. et al. Single-pot, solid-phase-enhanced sample preparation for proteomics experiments. Nat Protoc 14, 68–85, doi:10.1038/s41596-018-0082-x (2019).

50 Zhang, X. et al. Proteome-wide identification of ubiquitin interactions using UbIA-MS. Nat Protoc 13, 530–550, doi:10.1038/nprot.2017.147 (2018).

51 Lê, S., Josse, J. & Husson, F. FactoMineR: An R Package for Multivariate Analysis. Journal of Statistical Software 25, 1–18, doi:10.18637/jss.v025.i01 (2008).

52 Wu, T. et al. clusterProfiler 4.0: A universal enrichment tool for interpreting omics data. Innovation (Camb) 2, 100141, doi:10.1016/j.xinn.2021.100141 (2021).

53 Liao, Y., Smyth, G. K. & Shi, W. featureCounts: an efficient general purpose program for assigning sequence reads to genomic features. Bioinformatics 30, 923–930, doi:10.1093/bioinformatics/btt656 (2014).

54 Ritchie, M. E. et al. limma powers differential expression analyses for RNA-sequencing and microarray studies. Nucleic Acids Res 43, e47, doi:10.1093/nar/gkv007 (2015).

55 Granja, J. M. et al. ArchR is a scalable software package for integrative single-cell chromatin accessibility analysis. Nat Genet 53, 403–411, doi:10.1038/s41588-021-00790-6 (2021).

56. Garrison, E. & Marth, G. Haplotype-based variant detection from short-read sequencing. *arXiv e-prints*, arXiv:1207.3907, doi:10.48550/arXiv.1207.3907 (2012).

57 Yu, G., Wang, L. G., Han, Y. & He, Q. Y. clusterProfiler: an R package for comparing biological themes among gene clusters. OMICS 16, 284–287, doi:10.1089/omi.2011.0118 (2012).

